# The evolution of GC-biased gene conversion by means of natural selection

**DOI:** 10.1101/2024.06.21.600052

**Authors:** Augustin Clessin, Julien Joseph, Nicolas Lartillot

## Abstract

GC-biased gene conversion (gBGC) is a recombination-associated evolutionary process that biases the segregation ratio of AT:GC polymorphisms in the gametes of heterozygotes, in favour of GC alleles. This process is the major determinant of variation in base composition across the human genome and can be the cause of a substantial burden of GC deleterious alleles. While the importance of GC-biased gene conversion in molecular evolution is increasingly recognised, the reasons for its existence and its variation between species remain largely unknown. Using simulations and semi-analytical approximations, we investigated the evolution of gBGC as a quantitative trait evolving by mutation, drift and natural selection. We show that in a finite population where most mutations are deleterious, gBGC is under weak stabilising selection around a positive value that mainly depends on the intensity of the mutation bias and on the intensity of selective constraints exerted on the genome. Importantly, the levels of gBGC that evolve by natural selection do not minimize the load in the population, and even increase it substantially in regions of high recombination rate. Therefore, despite reducing the population’s fitness, levels of gBGC that are currently observed in humans could in fact have been (weakly) positively selected.

## 1 Introduction

In meiosis, during the repair of double strand breaks (DSBs), the single stranded DNA from the broken chromosome invades the homologue, such that the two form a double stranded DNA chimera (heteroduplex) of the two parental chromosomes. At this location, if the individual is heterozygous, there will be a mismatch (non Waston and Crick pairing). This mismatch can be resolved by repairing either parental allele with the other allele. This phenomenon therefore induces gene conversion (Winkler, 1930; Roman, 1985). In the late 80’s Brown and Jiricny (1987) found that in human and green monkey cells, DNA repair was biased towards GC alleles. Since then, direct and indirect evidence has revealed that repair biases operate on meiotic gene conversion events in a wide range of eukaryotes, leading to a biased transmission of GC alleles to offspring (Mancera et al., 2008; Duret and Galtier, 2009; Pessia et al., 2012; Smeds et al., 2016; Clément et al., 2017; Galtier et al., 2018; Boman et al., 2021). GC-biased gene conversion (gBGC) is therefore a special case of genome-wide non-Mendelian segregation where recombination and DNA repair machineries act as segregation distorters (Nagylaki, 1983; Bengtsson and Uyenoyama, 1990). Most methods that detect selection or infer demography from genetic data are based on the assumption of Mendelian segregation, and gBGC therefore confounds both selection and demography inference (Galtier and Duret, 2007; Ratnakumar et al., 2010; Kostka et al., 2012; Pouyet et al., 2017, 2018; Bolívar et al., 2019; Joseph, 2024). Moreover, it has been demonstrated, notably in humans and chickens, that gBGC is the major determinant of GC content variations along the genome (Galtier et al., 2001; Meunier and Duret, 2004; Webster et al., 2006).

Despite its major impact on genome evolution, the evolutionary origins of gBGC and the reasons for its maintenance remain quite uncertain. Bengtsson (1986) made the prediction that, if gene conversion could be biased against the most common class of mutations, it could provide an advantage by reducing the genetic load. GC *↦* AT mutations being the most common type in most species (Long et al., 2018), it has thus been naturally hypothesized that gBGC could have been selected as a correction mechanism that counteracts the almost universal mutational bias towards AT (Glémin, 2010). However, Glémin (2010) demonstrated that the levels of gBGC that should minimize the load are very weak compared to empirical values observed in regions of high recombination rate.

In fact, empirical studies so far are quite unanimous on a mostly deleterious effect of gBGC (Berglund et al., 2009; Galtier et al., 2009; Necşulea et al., 2011; Lachance and Tishkoff, 2014; Bolívar et al., 2016). Having a mechanism that seems to be mostly deleterious being so widespread in eukaryotes is therefore quite paradoxical. Interestingly, both in angiosperms and animals, studies observed a negative correlation between the transmission bias *b*, and effective population size (Clément et al., 2017; Galtier et al., 2018). Galtier et al. (2018) proposed that this pattern could be explained by a drift barrier hypothesis, whereby gBGC is a deleterious process which can be efficiently counter-selected only in species whose effective population size is high. However, as the way mutation, drift and selection affect the evolution of gBGC lacks theoretical expectations, this argument is verbal and requires theoretical validation. In this direction, Bengtsson and Uyenoyama (1990) investigated the evolution of a modifier of biased gene conversion (BGC) under different scenarios, and recovered that a positive value of BGC evolves naturally when mutation is biased. This result gives a stronger theoretical basis to the idea that gBGC could evolve as a consequence of an AT mutation bias. On the other hand, the study was conducted under the approximation of infinite population sizes and at a single strongly selected locus. As such, it does not provide an explanation for the variation in the strength of gBGC between species of different effective population sizes.

To tackle this question in a more realistic setting, we developed a model in which the intensity of gBGC evolves freely as a quantitative trait that affects the whole genome in a finite population. We confirm that, in the presence of a mutational bias towards AT, gBGC naturally evolves towards positive values (Bengtsson and Uyenoyama, 1990). As expected, the equilibrium value of the transmission bias towards GC depends both on the intensity of the mutational bias towards AT and on the magnitude of selective constraints exerted on the genome. Interestingly, we predict that the equilibrium value of the transmission bias correlates negatively with effective population size. Importantly, we show that even if gBGC leads to a higher deleterious burden at the population level, this does not mean that it is negatively selected, even in high *N*_*e*_ species. In the present model, high gBGC intensity results from the short-term advantage of converting AT deleterious alleles in heterozygotes, which leads to a higher deleterious burden in the population. Overall, by capturing the selective pressures acting on gBGC under empirically realistic conditions, this model provides insight into the role of natural selection in shaping the evolution of gBGC in eukaryotes.

## 2 Results

### Model summary

A model for the evolution of biased gene conversion was designed and implemented as a simulation program. The model is meant to represent a population of randomly-mating diploid individuals, of fixed size *N*, evolving under a typical nearly-neutral regime, that is, under purifying selection against deleterious mutations susceptible to occur over a broad (gamma-distributed) range of selective effects, from very weak to very strong (Ohta, 1992; Eyre-Walker and Keightley, 2007). Those mutations occur over a set of bi-allelic loci, with allelic states *W*, or Weak (corresponding to AT), and *S*, or Strong (corresponding to GC). The model assumes a mutation bias *λ*, which will be typically in favour of Weak alleles (so as to mimic the mutational bias in favour of AT seen across many eukaryotic species (Long et al., 2018)). Selection, on the other hand, is statistically balanced with respect to either *W* or *S*, in the sense that, for each locus, either *W* or *S* is randomly chosen to be the deleterious allele with probability 1*/*2.

On top of this nearly-neutral background, the model invokes a modifier locus, encoding an additive quantitative trait modulating biased gene conversion. Specifically, the locus determines the value of the conversion bias parameter *β*, which will play during meiotic recombination as follows: in addition to a unique cross-over uniformly chosen along the chromosome, a certain fraction of the genome undergoes gene conversion at rate *α* per nucleotide position. If a position somewhere in the genome is heterozygous and happens to undergo gene conversion, then the *W* allele is converted into the *S* allele with probability (1+*β*)*/*2, and conversely, the *S* allele is converted into the *W* allele with probability (1 *− β*)*/*2. As a result, the net strength of biased gene conversion, defined as the net excess of transmission of *S* alleles, relative to *W*, at a *WS* heterozygous position, is *b* = *αβ*.

The basal rate of gene conversion, *α*, is assumed to be fixed, possibly because of specific constraints related to the molecular mechanisms of meiosis. The conversion bias *β*, on the other hand, is allowed to evolve, by introducing mutant alleles at the modifier locus (at rate *w*) contributing a small shift, either positive or negative, in the value of *β*. As a result of this mutational input, biased gene conversion is susceptible to show variation among individuals. The whole question is then whether this genetically-encoded variation in gBGC is in turn subject to indirect selection, and whether this results in predictable patterns of evolution of gBGC in the long run.

### Biased gene conversion is under stabilizing selection

Typical trajectories of the population-mean of biased gene conversion (*b*) under the model are shown in Figure 1. Here, a mutational bias of *λ* = 3 is considered (bias in favour of Weak, or AT alleles), with a basal mutation rate of *u* = 10^*−*4^ (for *W* to *S* mutations), a population size of *N* = 1000, a genome consisting of *L* = 10000 selected loci, with a gamma distribution of selective effects of mean 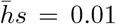 and shape 0.2. Two alternative settings are considered for the dominance effect of those mutations: either co-dominant (*h* = 0.5) or partially recessive (*h* = 0.1). In both cases, the modifier locus undergoes mutations at a rate of *w* = 10^*−*3^ per generation, with effect sizes of mean 0.1 on *β*. Finally, the basal gene conversion rate is equal to *α* = 0.1. Of note, these parameter values are not meant, at that stage, to match any specific empirical situation. Instead, the aim is to reveal the inner workings of the model, and how its output relates to the input parameter values.

**Figure 1:**
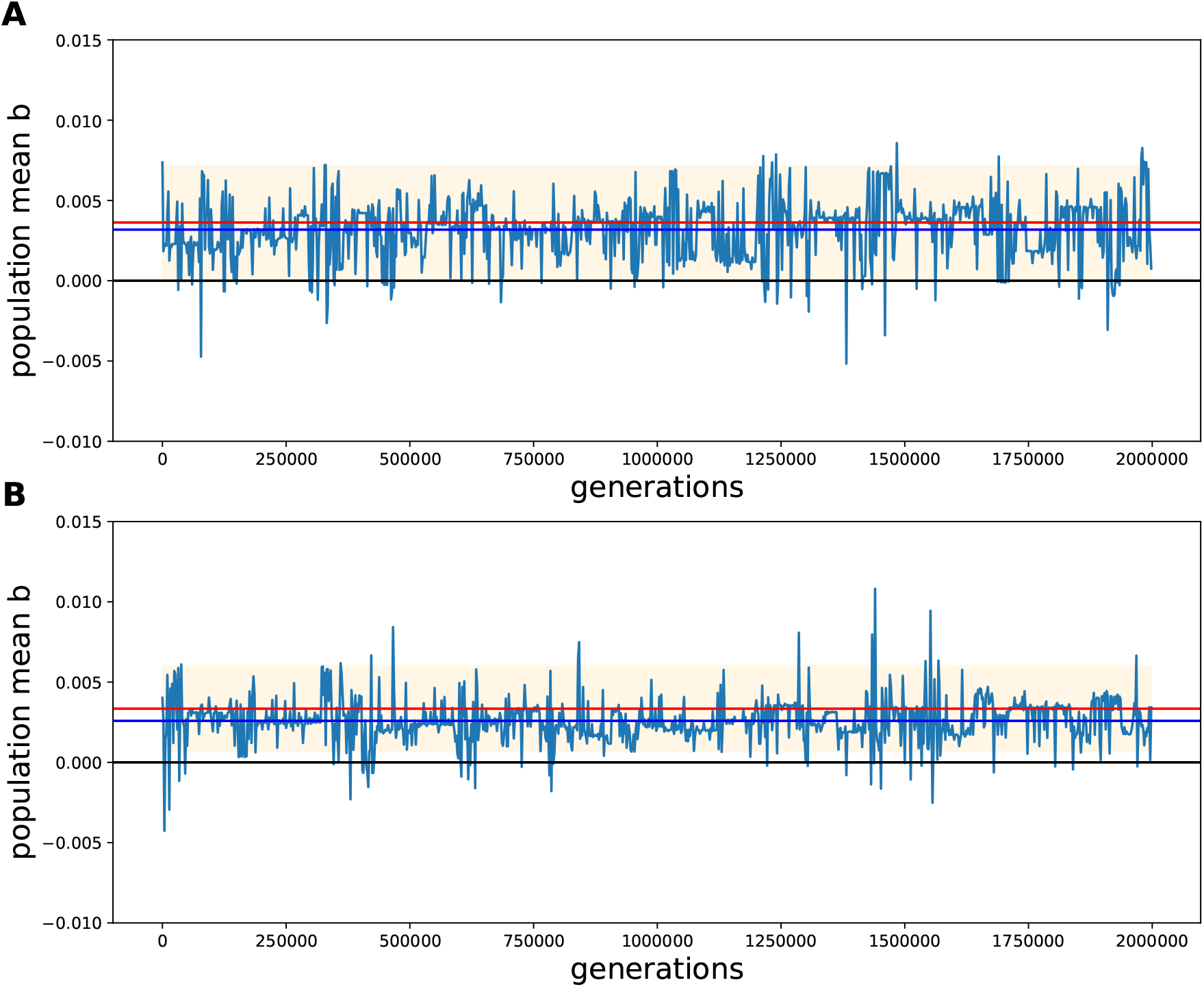
Evolution of the strength of gBGC (mean of *b* over the population) over the generations, for the co-dominant (A) and recessive (B) cases. Blue horizontal line: mean over the entire run; red horizontal line: equilibrium value predicted by the analytical approximation; shaded area: predicted equilibrium variance.

Running the model under these parameter values results in a population-level gBGC evolving towards positive value of *b*, reaching an evolutionary equilibrium with a long-term mean of the order of *b ≃* 0.003 (Figure 1). There is a substantial evolutionary variance, such that the population still spends about 5% of the time with negative values of *b*. Nevertheless, these experiments show that gBGC is susceptible to spontaneously evolve in favour of Strong (GC) alleles. They also more specifically suggest the existence of some form of stabilizing selection acting on gBGC, driving the population towards, and maintaining it around, an evolutionary equilibrium.

### The mutation-segregation tradeoff between AT- and GC-deleterious mutations

The observations gathered in the last section call for a deeper understanding of what drives the equilibrium value of *b*, and its variance. Given a mutation bias towards *W*, it seems relatively straightforward that a conversion mechanism playing blindly against *W* alleles during meiosis should be error-correcting on average and could therefore be selected (Bengtsson, 1986). What is perhaps less obvious is why selection induced on gBGC modifiers is stabilizing rather than extremal, resulting in an optimal value of *b*. The fundamental reason for this lies in the feedback of the evolution of gBGC on the segregation frequencies at the selected loci across the genome.

Consider a population initially devoid of gBGC. In this context, modifiers increasing gBGC are selectively favoured due to their error-correcting effect on deleterious polymorphisms, which are primarily towards W. Such modifiers will therefore invade. As a consequence, however, the population starts to live and reproduce under increasingly high levels of gBGC. This in turn changes the frequency at which *S* and *W* alleles segregate, increasing the frequency of *S* and decreasing the frequency of *W* alleles in the population. This shift in the segregation frequencies of deleterious alleles in favour of *S* tends to compensate for the mutation bias in favour of *W*. The balance between these two opposing effects, mutation versus segregation bias, is reached for an intermediate value of *b*.

This mutation-segregation tradeoff can be mathematically formalized under the assumption that gBGC evolves slowly and that most of the selection induced on gBGC is fundamentally contributed by selected loci that are not strongly linked to the modifier locus (these assumptions are discussed below). The detailed derivation is given in the Methods. Here, the main intuitions are presented and graphically illustrated.

The key is to express the mean selective effect induced on a modifier increasing the value of gBGC by an amount *δb*, in a population at equilibrium under a strength of gBGC equal to *b*. This induced selection is here more precisely defined as the difference between the mean fitness of the offspring of an individual bearing the modifier (and thus implementing a gBGC of strength *b* + *δb* in its meiosis) and the mean fitness of the offspring of an individual not bearing the modifier. For small *δb*, this difference is proportional to *δb* and can be written:

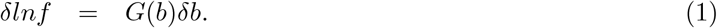

If *G*(*b*) is positive, then modifiers increasing *b* will be favoured, and conversely if *G*(*b*) is negative. Considering *W* and *S* alleles separately, *G*(*b*) can be expressed as the difference between the net gain upon converting *W* -deleterious alleles *G*_*W*_ (*b*), and the cost of converting *S*-deleterious alleles, *G*_*S*_(*b*):

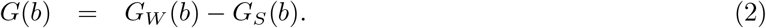

Both terms are positive, and the sign of *G*(*b*) will thus be determined by which of these two contributions, gain or cost, is largest.

Since the selection induced on gBGC is contributed by the entire genome, both *G*_*W*_ (*b*) and *G*_*S*_(*b*) can be expressed as averages over the distribution of selective effects of the mean selective impact of gene conversion events, scaled by the number of positions under selection, which is *L/*2 for both cases:

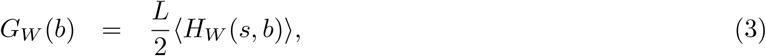

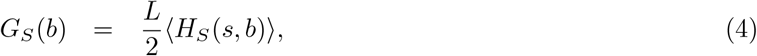

Here, *H*_*W*_ (*s, b*) and *H*_*S*_(*s, b*) denote the mean selective impact of gene conversion events at loci with selection coefficient *s*, at equilibrium under a gBGC equal to *b*. The angle brackets stand for an expectation over the gamma distribution of selective effects.

Finally, in order to account for the stochastic fluctuations in the segregation frequencies of selected loci, the functions *H*_*S*_(*s, b*) and *H*_*W*_ (*s, b*) are themselves expectations over the frequency distribution for *W* and *S* alleles, of the expected selective differences contributed in the offspring by conversion events occurring during meiosis on the selected positions that happen to be heterozygous in a typical individual. Thus, taking the case of *W* -deleterious alleles, let *x* denote the frequency at which the allele segregates in the population.

Under random mating, an individual will be heterozgous for this allele with probability

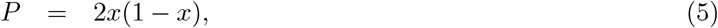

in which case, accounting for all possible genotypes for the other parent, the mean gain induced by a conversion event at that position in the offspring will be equal to (see methods):

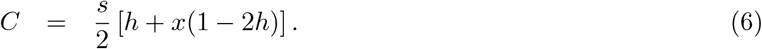

At mutation-selection-conversion balance, *x* is a random variable drawn from an equilibrium frequency distributions noted 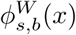, and thus, overall, the mean gain will amount to:

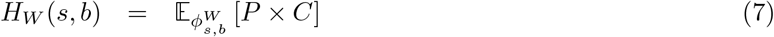

The same derivation can be conducted in the case of *S*-deleterious loci. For bi-allelic loci, the equilibrium distributions 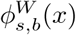 and 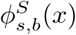, for both *W* and *S* loci, can be explicitly written, up to a normalization constant, such that expectations over these distributions can be computed numerically (see methods).

### Predicting the equilibrium mean and variance strength of gBGC

The value of *G*(*b*) = *G*_*W*_ (*b*) *− G*_*S*_(*b*) can be plotted as a function of the population-level *b* (Figure 2). This function is decreasing, crossing 0 at an intermediate, positive value of *b*^*∗*^. Numerically solving for the value *b*^*∗*^ such that *G*(*b*^*∗*^) = 0 gives *b*^*∗*^ = 0.0036 in the co-dominant case, and *b*^*∗*^ = 0.0033, which is close to the mean value observed in the simulation (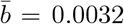 and 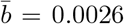, respectively). The numerical and simulation-based estimates are both represented as a blue and red lines, respectively, in Figure 1.

**Figure 2:**
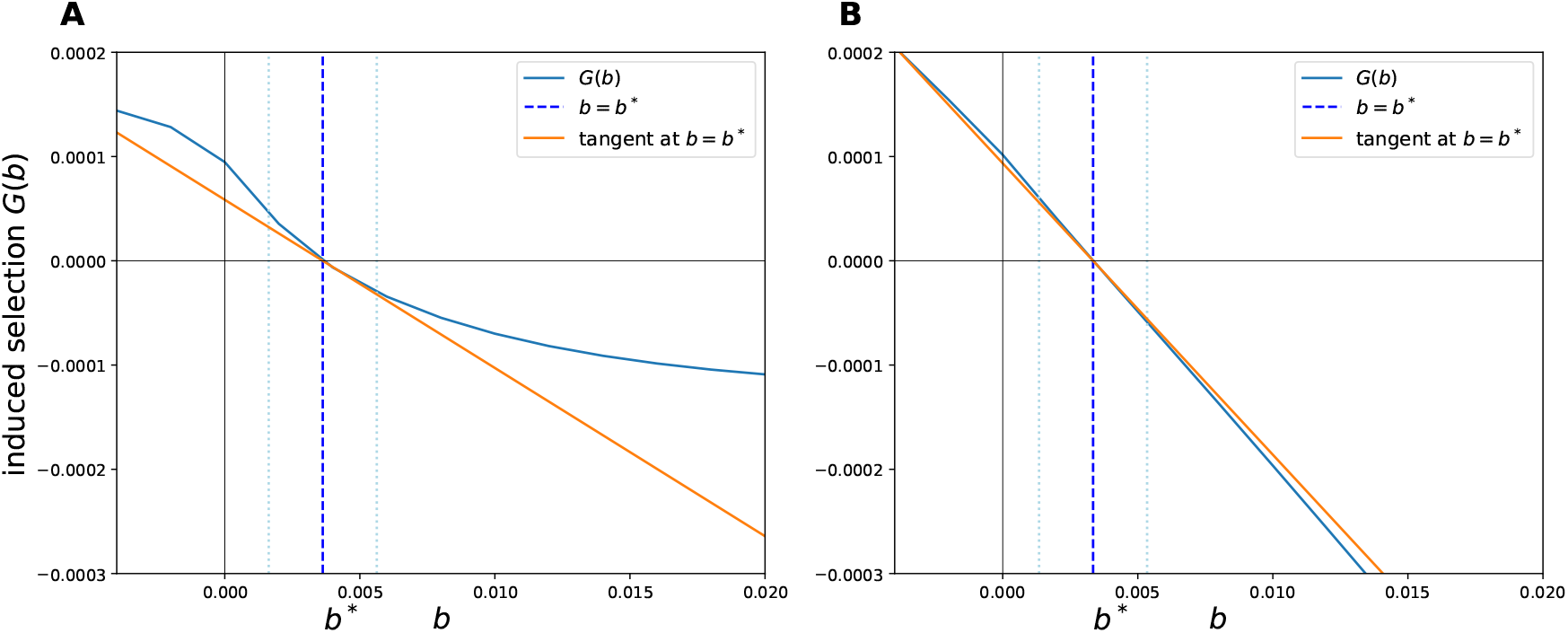
Strength of selection induced on gBGC modifiers, *G*(*b*), as a function of *b* (blue curve), under the co-dominant (A) and recessive (B) settings. Dark blue dotted vertical line: numerically estimated value of *b*^*∗*^, for which *G*(*b*) = 0; orange line: numerically estimated tangent at *b*^*∗*^; light blue dotted vertical lines: predicted standard deviation around *b*^*∗*^.

A rough quantitative estimate of the evolutionary variance can also be obtained, based on the slope *γ* of the tangent to the curve at *b*^*∗*^ (Figure 2A). Specifically, the equilibrium evolutionary variance is predicted to be approximately equal to 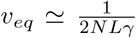 (reported as a shaded area on Figure 1). Of note, the selective response shows a steeper slope at the equilibrium point in the recessive case, resulting in a smaller predicted evolutionary variance than in the co-dominant case.

### The drivers of gBGC

The behaviour of the simulation model, along with the analytical approximation just introduced, were further investigated by plotting the predicted equilibrium value of the strength of gBGC, *b*^*∗*^, as a function of several key parameters (mutation bias, mean strength of purifying selection, number of positions under selection and mutation rate). The case of the response of *b*^*∗*^ to changes in effective population size is examined further below.

Not surprisingly, the mean equilibrium strength of gBGC is directly related to the strength of the mutational bias (Figure 3A). Owing the symmetry of the problem, running the model with *λ <* 1, i.e. under a mutational bias in favour of the Strong alleles results in a population evolving towards a mean conversion bias in favour of Weak (left side of Figure 3A). The mean equilibrium strength of biased gene conversion is also directly influenced by the mean strength of the purifying selection acting over the genome (Figure 3B), thus clearly indicating that its evolutionary dynamics is a direct consequence of the selective effects induced by converting non-neutral polymorphisms in the germ-line. The mean equilibrium value is insensitive to the number *L* of selected loci, but its evolutionary variance, on the other hand, is affected, showing a clear decreasing trend with *L*, which corresponds to the scaling in 1*/L* predicted by the analytical approximation (Figure 3C).

**Figure 3:**
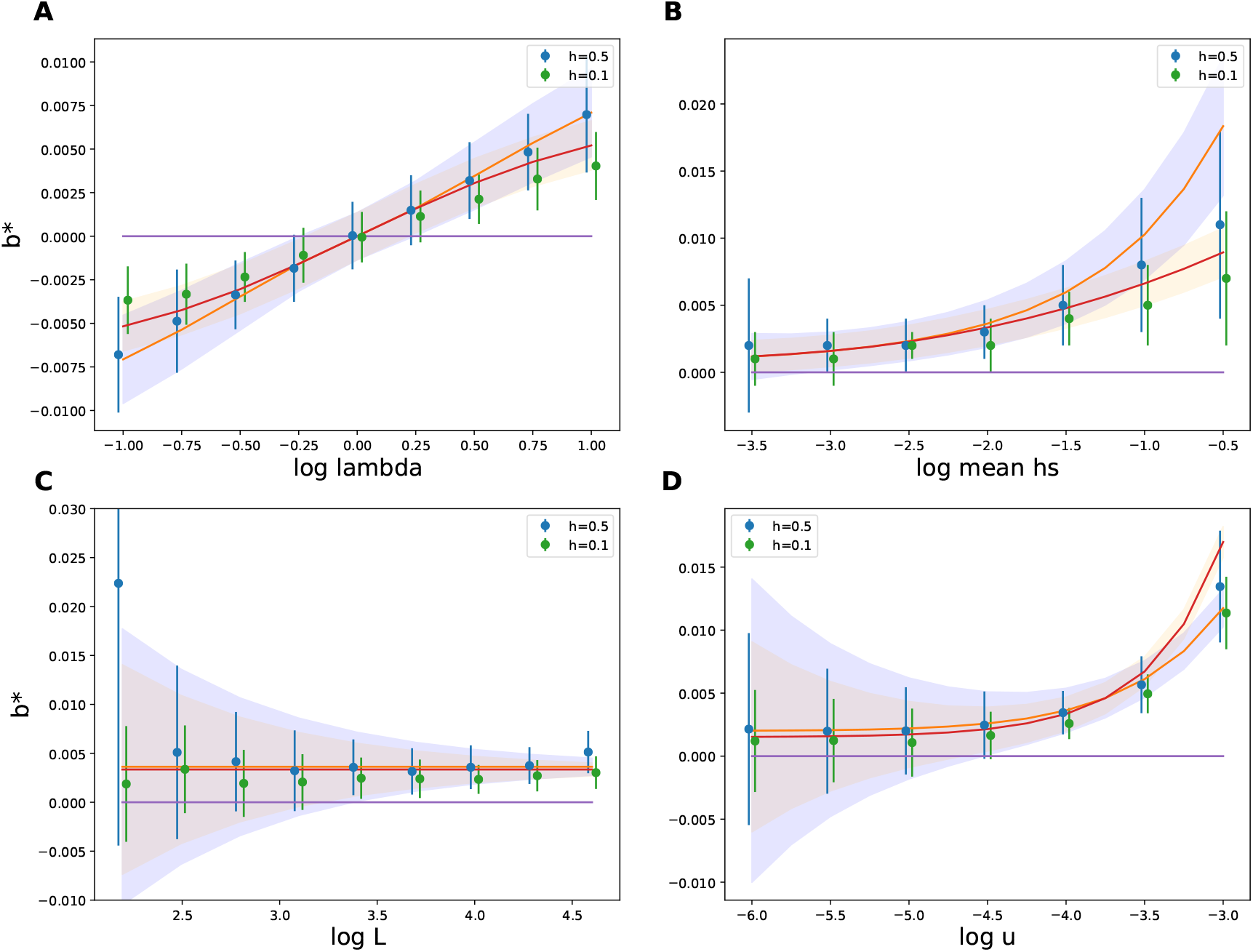
Mean equilibrium *b*^*∗*^ and standard deviation, as a function of mutation bias *λ* (A), mean selective effect 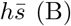, number of selected positions in the genome *L* (C) and mutation rate *u* (D), under the co-dominant (blue) and the recessive (orange) case, obtained by simulations (dots and associated vertical bars) and predicted by the analytical approximation (curve and associated shaded area), under a mutation rate of *w* = 10^*−*4^ (*Nw* = 0.1).

Finally, the strength of gBGC responds very weakly to the mutation rate, except for very high mutation rates (4*Nu >>* 1), in which case it shows a sharp increase (Figure 3D). For low 4*Nu*, not so much the mean than the evolutionary variance of gBGC is impacted by the mutation rate, with larger variances being observed under lower mutation rates. In this respect, the response of the model to variation in *u* is not unlike its response to variation in *L* (Figure 3C). This similar behaviour can be understood by noting that any indirect selective effect acting on the modifier locus can only be mediated by heterozygous positions. Thus, the strength of induced selection will be directly determined, not just to *L*, but more fundamentally, by the mean number of selected positions at which a typical individual is heterozygous. The mean heterozygosity in the population is in turn directly impacted by the mutation rate, and this, under most selective regimes.

The analytical approximation (plain lines in Figure 3) is globally in good agreement with the simulation results (filled circles), except for large 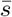 or large *u*, where it underestimates the equilibrium strength of gBGC.

However, these corresponds to regimes where the diffusive approximation used for deriving the analytical predictions is not valid, owing to a large variance in genome-wide log-fitness between individuals. In practice, these regimes are far from empirical reasonable conditions.

Finally, across all scaling experiments shown in Figure 3, the stabilizing selection induced on *b* appears to be globally tighter in the partially recessive case, for which both the response of the equilibrium value of *b* to changes in parameter values and the equilibrium variance are less pronounced than in the co-dominant case.

### Which class of mutations contribute to stabilizing selection on gBGC ?

As the mean fitness effect of deleterious mutations is a key parameter for the evolution of intermediate levels of gBGC, it appears probable that under a distribution of fitness effects (DFE), not all mutations contribute equally to it. To further investigate this point, the analytical approximation was recruited to examine how the mean frequency at which *W* -deleterious alleles (Figure 4A&B) and *S*-deleterious alleles (Figure 4C&D) segregate in a population is modulated by slight variations in *b* (the dotted, plain, and dashed lines correspond to increasingly larger values of *b*). The bottom panels show the corresponding expected fitness gain *H*_*W*_ (*s, b*) incurred by converting *W* -deleterious alleles (blue curves, above 0), and the expected fitness cost *H*_*S*_(*s, b*) incurred by converting *S*-deleterious alleles (red curves, below 0), both weighted by the distribution of selective effects (DFE). These are plotted as functions of *s*, for 3 different values of *b*. Weighting *H* by the DFE gives a better sense of the relative contributions of mutations with different selection coefficients to the total cost and gain. Also, with this weighting, averaging *H*_*W*_ and *H*_*S*_ over the DFE simply amounts to computing the area under the two curves, which thus directly correspond to *G*_*W*_ (*b*) and *G*_*S*_(*b*), respectively. The parameter values used for Figure 4 correspond to the simulation trajectory displayed in Figure 1, for the co-dominant and recessive cases.

**Figure 4:**
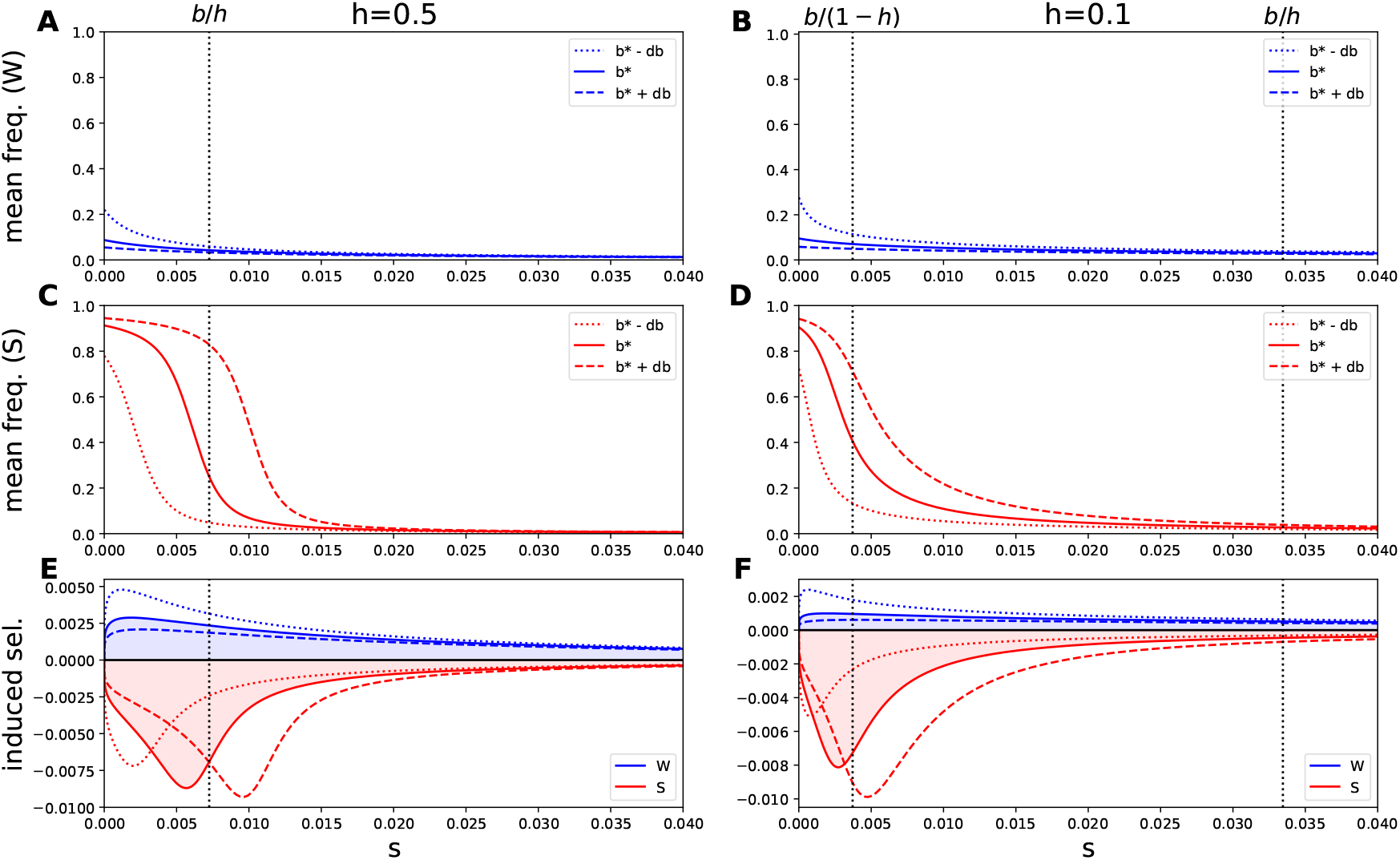
Mean segregation frequency of *W* alleles (A&B), *S* alleles (C&D), and induced selection (E&F), as a function of *s*, under the co-dominant (A,C&E) and recessive (B,D&F) settings, for different values of *b*: plain lines correspond to the equilibrium value of *b*, dashed lines to a slightly increased *b*, and dotted lines to a slightly decreased *b*. Blue lines correspond to W alleles, while red lines correspond to S alleles.

As *b* increases, *W* alleles segregate at a lower frequency (Figure 4A&B) and *S* alleles at a higher frequency (Figure 4C&D). Correlatively, the expected gain contributed by converting *W* alleles (Figure 4E&F, blue curves) decreases, and the cost contributed by converting *S* alleles (red curves) increases with population-level *b*. The intermediate value of *b* used in Figure 4 is precisely the one for which the areas under the two curves in panel C are equal (shaded areas in blue and red) – it is thus the predicted evolutionary equilibrium (of note, the areas under the two curves may not look equal to each other on the figure, in particular in the recessive case, but this is only because the two curves extend much further to the right than is shown).

Importantly, the way the two compartments, Weak and Strong, react to changes in population-level *b* is very different. On one side, *W* -deleterious polymorphisms are only moderately affected, and this, mostly in the range of small selective effects. In contrast *S*-deleterious polymorphisms are strongly affected. More specifically, increasing *b* leads to a surge in the segregation of *S*-deleterious mutations of intermediate strength, for which gBGC and selection are of the same order of magnitude. This surge translates into a peak in the expected cost (red curve, bottom panel), whose area increases with *b*. Translating these observations in terms of the net selection acting on gBGC, the fitness advantage is mostly contributed by converting *W* strongly-deleterious mutations, and is essentially a constant. The fitness cost, on the other hand, is mostly contributed by *S* mutations of selective effects of the order of *b*. This fitness cost varies strongly with *b* and is the main factor responsible for modulating the selection induced on gBGC modifiers, as a function of the population *b*.

Of note, the exact patterns differ between the co-dominant and the recessive cases. In the co-dominant case, the peak in the conversion cost is around *s ≃ b/h*, the value for which gBGC and selection exactly compensate each other. In the recessive case, the region that contributes to the increased cost when gBGC increases is between *b/*(1 *− h*) *≤ s ≤ b/h*, or equivalently *hs ≤ b ≤* (1 *− h*)*s*. This is the range for which the strength of gBGC is stronger than selection against the heterozygote but weaker than selection against the homozygote for the deleterious mutant. As a result, these GC-deleterious polymorphisms tend to segregate at intermediate frequencies, as if they were over-dominant (i.e. advantageous when in one copy, deleterious when in two copies), resulting in a higher fraction of heterozygotes in the population, and thus a substantial cost against gBGC (Glémin, 2010). This can explain why the strength of selection around the equilibrium value of *b* is higher in the partly recessive case.

### gBGC is partially buffered against changes in population size

A somewhat paradoxical consequence of gBGC is the extreme sensitivity of equilibrium base composition to even mild variation in its population-scaled intensity *B* = 4*Nb* (Eyre-Walker, 1999). Quantitatively, the neutral equilibrium GC/AT composition ratio scaling exponentially with *B*, which can quickly lead to very large GC content even for moderate increase in *N*. For instance, based on the current estimate of *b* in humans, increasing effective population size by a factor 10 would imply a long-term neutral equilibrium GC content greater than 99% in the 10% most highly recombining fraction of the genome. How to explain, then, that gBGC does not more often lead to diverging base composition across species?

Implicit in the argument just exposed is that the strength of gBGC is fixed, while population size varies, or at least, that there is no internal mechanism for tuning the raw intensity of gBGC (*b*) depending on effective population size (*N*), so as to somehow guarantee that *B* = 4*Nb* never becomes too large. Yet, if gBGC is under stabilizing selection, this raises the possibility for such an internal mechanism to spontaneously emerge. This fundamentally depends on how the evolutionary optimum *b*^*∗*^ scales with population size.

To examine this point, the optimal value *b*^*∗*^ predicted by the model was computed (using the semi-analytical approximation) over a broad range of values of *N* between 10^2^ and 10^6^. For this experiment, a mutation rate of *u* = 10^*−*8^ was assumed (for *S ↦ W* mutations), and a bias of *λ* = 2. Both the co-dominant case (*h* = 0.5) and the partially recessive case (*h* = 0.1) were considered.

In all cases (Figure 5), whether co-dominant or recessive, *b*^*∗*^ decreases with *N*. The trend is moderate in the co-dominant case but more pronounced in the recessive case. In both cases, the decrease is less than linear, such that *B* = 4*Nb* still increases as a function of *N*. This increase is quite substantial in the co-dominant case, with *B* reaching values above 10 for population sizes of *N* = 10^4^ and above 100 for *N >* 3.10^5^. In the recessive case, on the other hand, *B* is much less responsive to changes in population size, ranging from *B ≃* 3 for *N* = 10^2^ up to *B ≃* 15 for *N* = 10^6^ – barely a 5-fold increase over 4 orders of magnitude for *N*.

**Figure 5:**
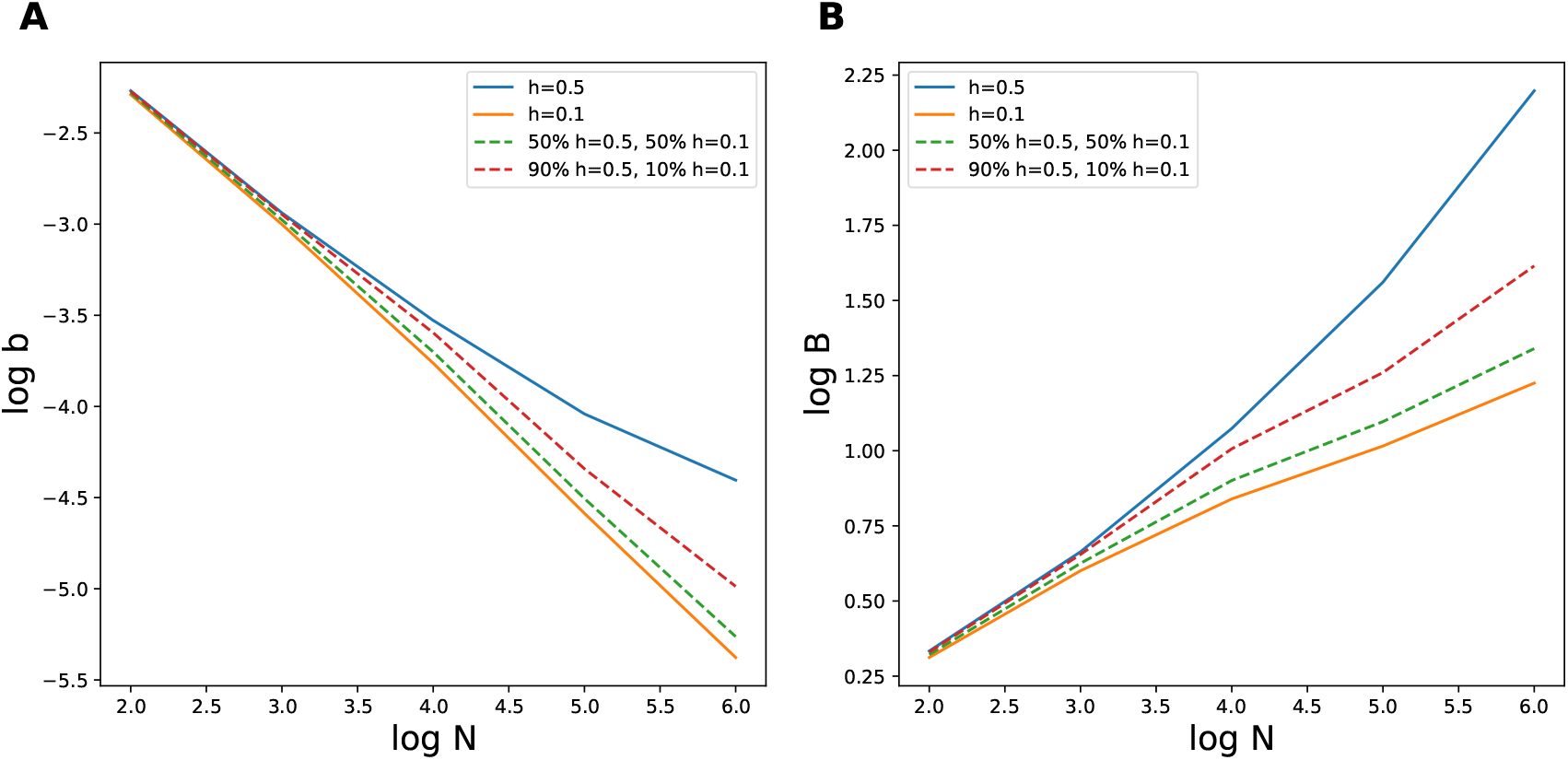
Scaling of *b*^*∗*^ (A) and *B*^*∗*^ = 4*Nb*^*∗*^ (B) as a function of *N*, under the co-dominant (*h* = 0.5) and recessive (*h* = 0.1) settings (plain curves), or assuming a mixture of co-dominant and recessive mutations (dashed curves).

Interestingly, a mixture of 50% co-dominant and 50% partially recessive (*h* = 0.1) essentially behaves like the pure partially recessive case (all mutations with *h* = 0.1). Even a small proportion of 10% of partially recessive positions, mixed with 90% of co-dominant positions, shows substantially more stable levels of gBGC as a function of *N* (Figure 5, dashed lines). Recessive mutations thus appear to represent an efficient buffer against changes in population-scaled gBGC induced by changes in population size.

The fundamental reason why *b*^*∗*^ decreases with *N* can be understood by examining the structure of the induced selective response (Figure 6). As mentioned above, the mutation-segregation balance essentially takes the form of a tradeoff between, on one side, a net error-correcting effect on strongly deleterious mutations (more often deleterious towards *W* than towards *S*) and, on the other side, a conversion load mostly contributed by *S*-deleterious mutations with selection coefficients of the order of *b*. The first component, being in the strong selection regime, is essentially insensitive to *N* (Figure 6E&F, blue curves). The second component, on the other hand, precisely because of the compensation between gBGC and selection, is effectively in a regime dominated by drift, and thus, in many respects, has an evolutionary dynamics resembling nearly-neutral evolution. As such, its mean heterozygosity is strongly influenced by changes in effective population size, and more precisely, will tend to increase with *N*. Since biased gene conversion is in direct proportion to the amount of heterozygosity, the conversion cost itself will also increase with *N* (Figure 6E&F, red curves). Altogether, GC-deleterious mutations with selective effects of the order of *b* are efficiently mobilized (i.e. contribute more to standing variation) upon an increase in *N* and thus represent a key force buffering *B*^*∗*^ against changes in population size.

**Figure 6:**
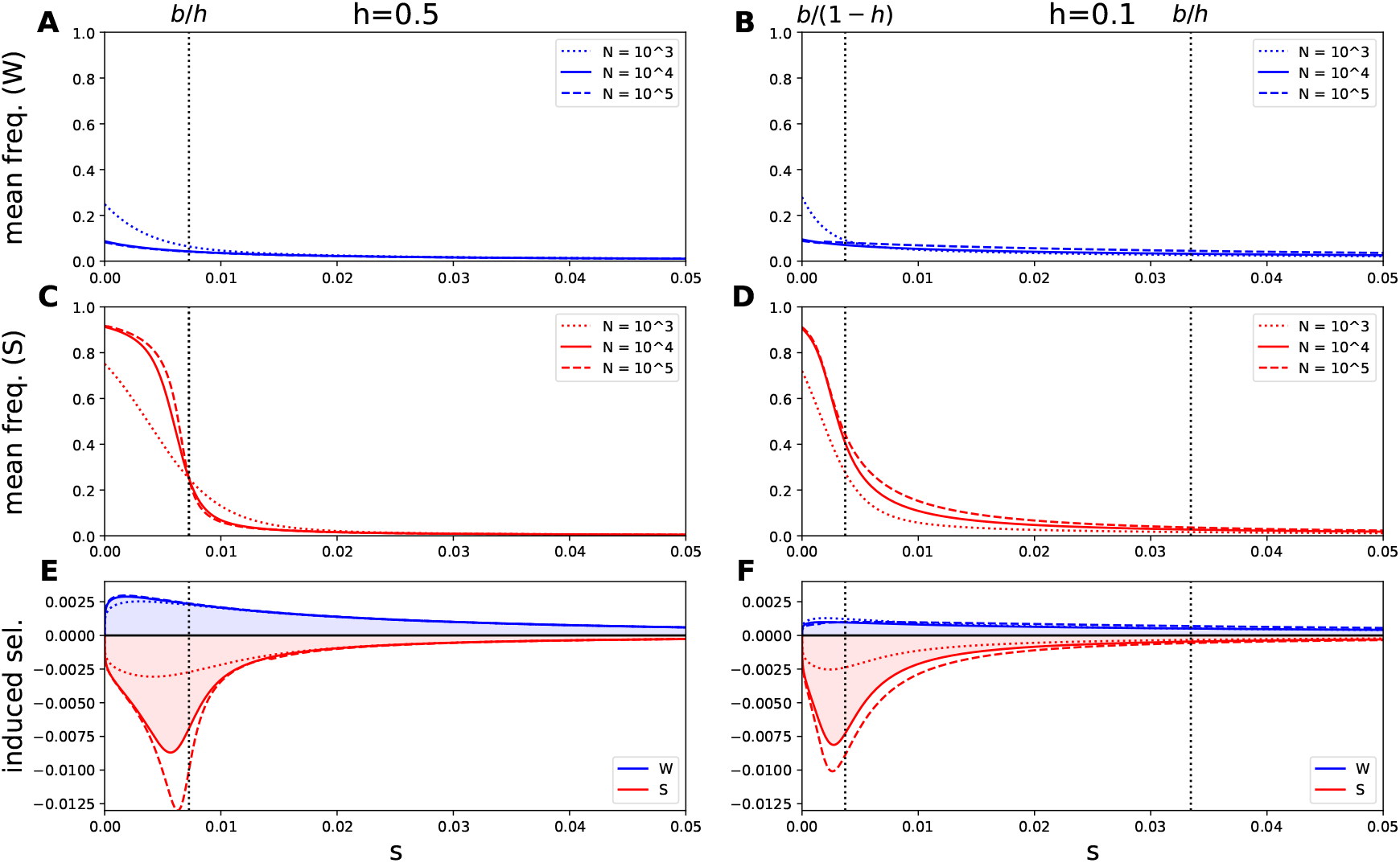
Mean segregation frequency of *W* alleles (A&B), *S* alleles (C&D), and induced selection (E&F), as a function of *s*, under the co-dominant (A,C&E) and recessive (B,D&F) settings, for different values of *N* : plain lines correspond to the equilibrium value of *b*, dashed lines to a slightly increased *N*, and dotted lines to a slightly decreased *N*. Blue lines correspond to W alleles, while red lines correspond to S alleles.

Of note, and as already explored above (Figure 4), in the co-dominant case (Figure 6, A,C&E), the range of GC-deleterious mutations that are mobilized consists of a relatively narrow peak around *b/h*. In contrast, in the recessive case, a good fraction of the range comprised between *b/*(1 *− h*) and *b/h* (the two dotted vertical lines on Figure 6B,D&F), corresponding to the co-dominant regime, is mobilized, thus contributing a much more responsive buffer against changes in *N* – which can easily dominate the overall response even if recessive mutations represent a minority of the total standing variation, as observed above (Figure 6, dashed lines).

### gBGC and the genetic load

gBGC is often depicted as a force that interferes with selection, and that causes a significant deleterious burden (Galtier and Duret, 2007; Berglund et al., 2009; Galtier et al., 2009; Necşulea et al., 2011). On the other hand, our results reflect those of previous studies showing that BGC confers a significant fitness advantage by correcting the most common class of mutations (here *S* ↦ *W*) (Bengtsson and Uyenoyama, 1990). But as pointed out by Glémin (2010), the levels of gBGC that evolve naturally are not necessarily the ones that minimize the average genetic load of a population. The average genetic load of a population can be decomposed into the load of W deleterious alleles:

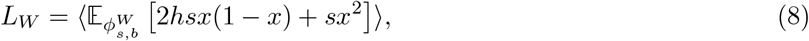

where *x* is a random variable drawn from the equilibrium frequency distribution of *W* alleles 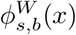, and that of S deleterious alleles:

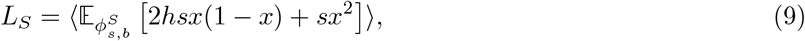

where *x* is a random variable drawn from the equilibrium frequency distribution of *S* alleles 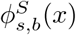 In both cases, and as above, the inner expectation is on *x* and the outer expectation (angle brackets) is on *s* (drawn from the gamma DFE).

Using a population size of *N* = 10, 000, a mutation rate of *u* = 10^*−*8^ and a mean selection coefficient *hs* = 0.01, we computed the average genetic load as a function of *b* for *h* = 0.5 and *h* = 0.1 (Figure 7A&B). The load is minimized for a very small value of *b* compared to the one that naturally evolves. The level of gBGC that minimizes the average genetic load corresponds to the level that equalizes the frequencies of *W* and *S* alleles, leading to an average GC content of 0.5 (Figure 7C&D). It is worth noting that when the average frequencies of *W* and *S* alleles are equal, it does not mean that they are distributed evenly in heterozygotes. In fact, *W* deleterious alleles are more often heterozygous (Figure 7 E&F), because they are numerous due to a high *S* ↦ *W* mutation rate, but at low frequency because of gBGC. Conversely, *S* deleterious alleles are less numerous due to a high *S* ↦ *W* mutation rate, but more often at high frequency because of gBGC and thus more often homozygous. Therefore, when the mean load in the population is minimal, there still is an individual advantage to convert *W* deleterious alleles more often for heterozygotes.

**Figure 7:**
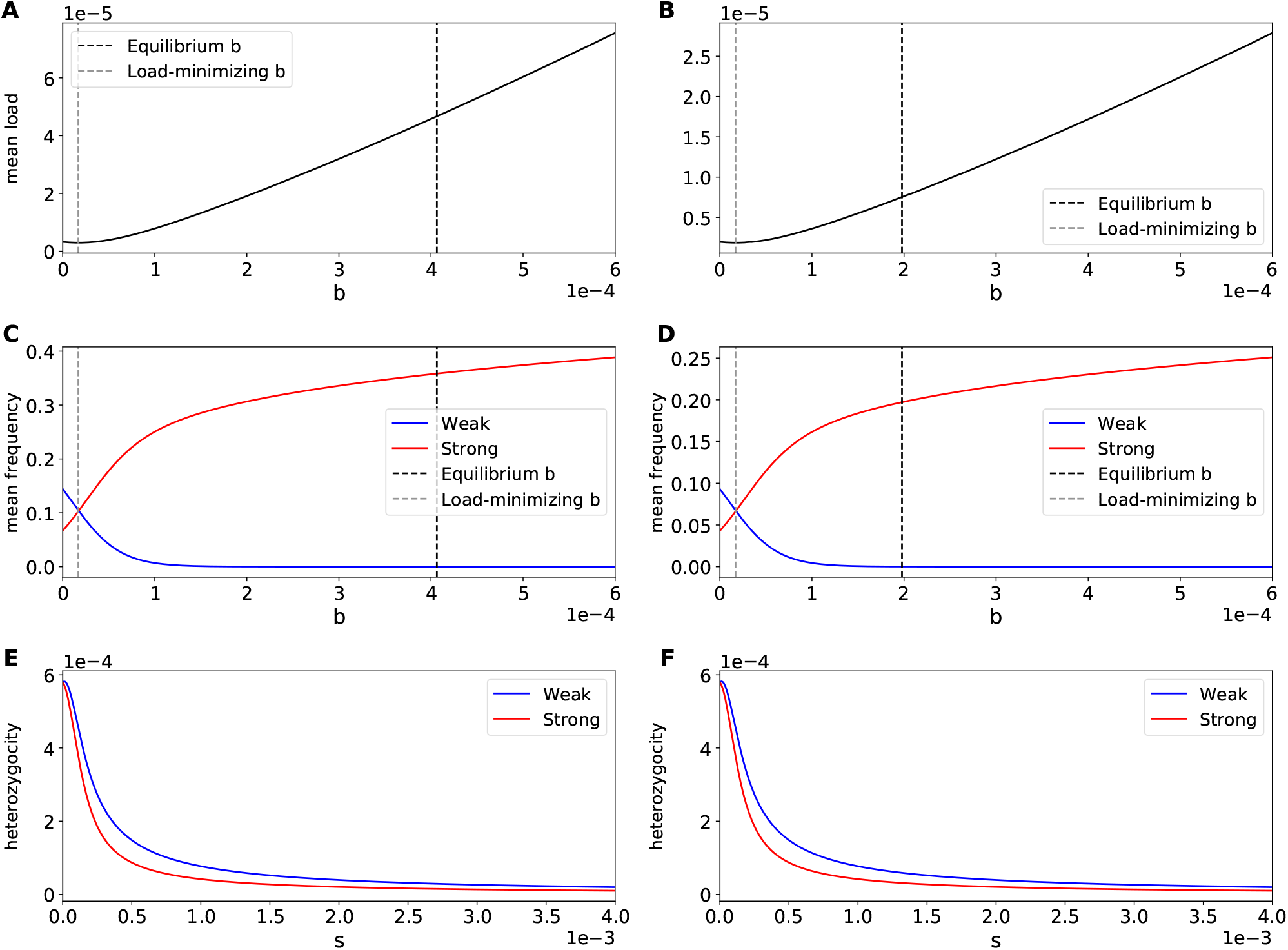
A&B: average deleterious load in a population as a function of *b*. The black lines shows the value equilibrium value of *b* and the grey line shows the value of *b* that minimizes the average load. C&D: frequency of W and S alleles as a function of *b*. E&F: heterozygosity for *W* ans *S* alleles as a function of their deleterious effect *s* under the value of *b* that minimizes the load. A,C&E: *h* = 0.5. B,D&F: *h* = 0.1

### Empirical calibration

The modeling work presented thus far suggests that biased gene conversion in favour of GC can in principle evolve as a consequence of the mutation bias towards AT and that its intensity can also be modulated in an adaptive manner as a function of key parameters, in particular effective population size. An important question that remains is whether the model provides quantitatively reasonable predictions when confronted to current empirical knowledge about the strength of gBGC in various species.

Humans and the mouse represent good cases to consider. To account for the substantial heterogeneity in recombination rates across the genome of these two species, the theoretical calculations were adapted so as to allow for a mixture of gBGC strength, whose mean is under the control of the modifier. Quantitatively, based on the estimate that about 90% of the recombination is concentrated in about 10% of the genome (Smagulova et al., 2011; Pratto et al., 2014), we assume that 10% of the genome experiences a 100 times stronger gBGC than the remaining 90%. For the other parameters, assuming that *∼*10% of the genome is under selection (Rands et al., 2014), for a genome of total size 3 Gb, this gives *L* = 3.10^8^ positions. In humans, the mutation rate is *u* = 3.10^*−*8^ per base pair and per generation. In the mouse, the mutation rate is a bit lower *u* = 10^*−*8^. Here, only *u* = 3.10^*−*8^ is considered. The mutation bias is in both cases of the order of *λ* = 2, the value used here. Current estimates of the DFE suggest a shape parameter *a* between 0.15 and 0.25 (Eyre-Walker et al., 2006; Castellano et al., 2019). Here, we used three values for *a*: 0.1, 0.2 and 0.3. The mean selection coefficient under this DFE is difficult to estimate. In humans, recent estimates are of the order of 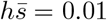 to 0.05, both of which were tried in what follows. Finally, the co-dominant and partially recessive cases are considered, as well as the 50:50 and 90:10 mixtures of these two dominance regimes, with population sizes varying from *N* = 10^4^ to *N* = 10^6^, so as to cover most of the range of what can be expected more generally in mammals.

The estimates of *B*^*∗*^ = 4*Nb*^*∗*^ (mean over the genome) under these parameter values are reported in Table 1. Under co-dominant selection, the predicted values for *B*^*∗*^ range from 1.5 to 124, showing quite some sensitivity to effective population size, mean selection strength across the genome, and shape of the DFE (Table 1 and Table S2&3). In contrast, assuming partially recessive mutations returns a much narrower range of estimates, from 0.8 to 6. Fitting the model assuming a mix of co-dominant and recessive mutations (last rows of Table 1) suggests that a moderate fraction of recessive mutations is sufficient to make *B* less responsive to changes in *N*.

**Table 1:**
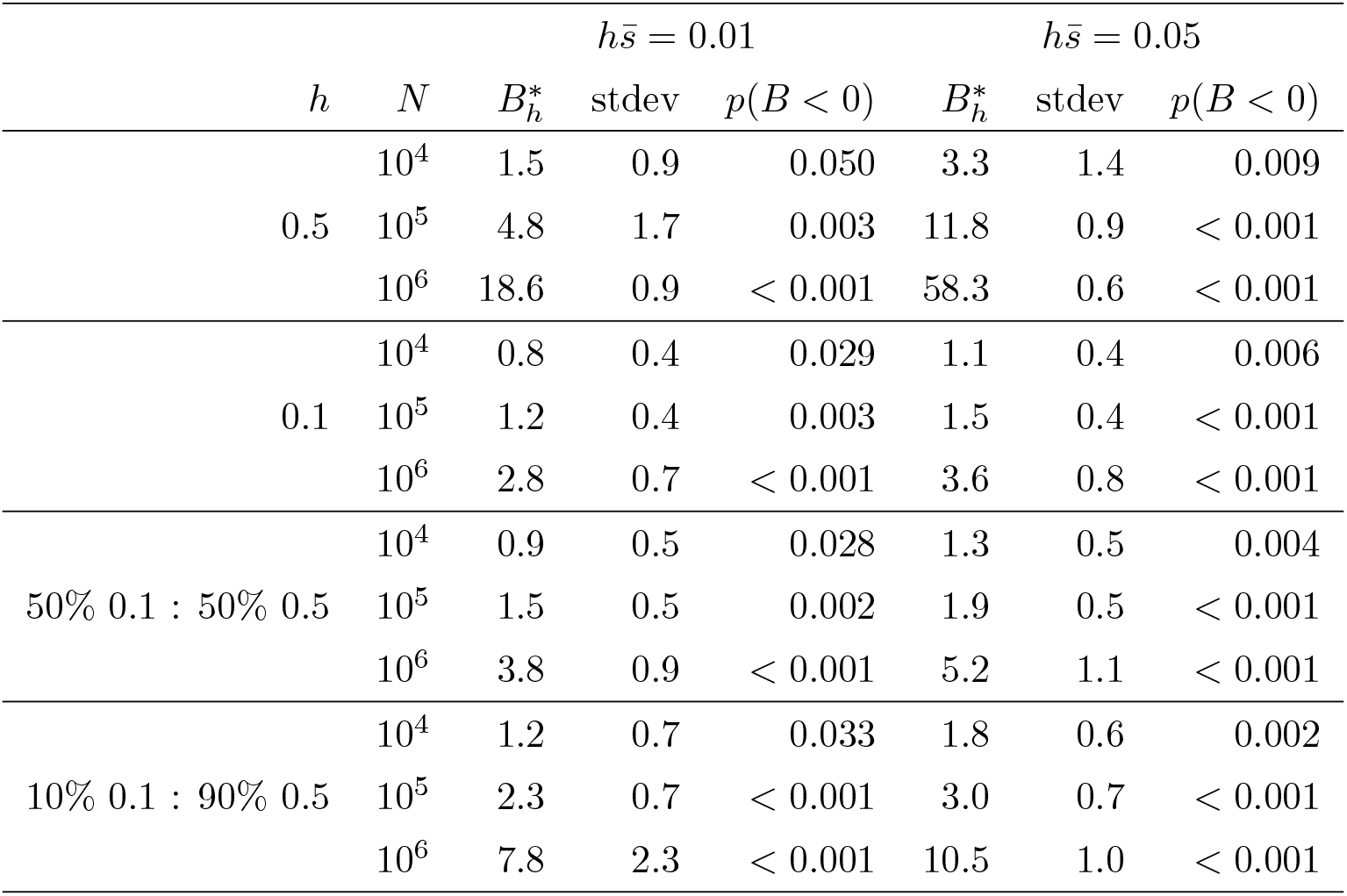
Numerical estimates of scaled intensity of gBGC *B*^*∗*^ = 4*Nb*^*∗*^ (mean over the genome), equilibrium standard deviation and probability of a negative gBGC, for different parameter values for *N*, *h*,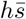.

Empirical estimates of gBGC in Humans are of the order of *B* = 0.3 for the genome-wide mean, and around *B* = 5.2 to 6.5 in the top 20% regions of high recombination (Duret and Arndt, 2008). The theoretical predictions (Table 1) are globally higher than these empirical estimates, although they get reasonably close to them (predicted *B* of the order of 1) for the lower values of 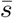 or assuming the presence of recessive mutations. This, together with the rather extreme results obtained for the largest population size under the co-dominant case, suggest that recessive mutations may play a role in buffering gBGC.

Finally, the predicted evolutionary variance is small, although not negligible for low population size, for which *b*^*∗*^ is predicted to be negative around 1 to 5% of the time. This suggests that induced selection on gBGC may not always be sufficiently powerful to guarantee a bias towards Strong over the whole range of molecular evolutionary regimes susceptible to be observed across mammals – although, even then, it may still represent a sufficiently strong selective force preventing gBGC to become unreasonably large.

## 3 Discussion

In this study, we developed a model to characterize the evolution of gBGC by means of natural selection. We first showed that, in the presence of a mutation bias towards AT, gBGC was under relatively weak stabilizing selection around a positive value of the transmission bias, in agreement with a previous study (Bengtsson and Uyenoyama, 1990). The equilibrium value of the transmission bias (*b*^*∗*^) corresponds to the one that equalizes the fitness gain of converting strongly deleterious AT mutations in heterozygotes and the fitness cost of transmitting slightly deleterious GC mutations to offsprings. This balance depends both of the strength of the mutation bias towards AT, but also on the mean fitness and dominance effects of deleterious mutations. When even few deleterious mutations are recessive, the cost of transmitting slightly deleterious GC alleles becomes quickly higher, and *b*^*∗*^ decreases. Importantly, *b*^*∗*^ is negatively correlated to effective population size. In fact, the fitness gain of correcting strongly deleterious AT mutations is essentially independent of effective population size, while the cost of transmitting slightly deleterious GC alleles increases quickly with it. This could contribute to the absence, or the weak positive correlation between the population-scaled gBGC coefficient (*B* = 4*N*_*e*_*b*) and effective population size (*N*_*e*_) reported in several clades of eukaryotes (Lartillot, 2013; Clément et al., 2017; Galtier et al., 2018; Galtier, 2021; Boman et al., 2021).

### Decoupling the short- and long-term effect of gBGC

gBGC is often described as an evolutionary force that antagonizes natural selection. It has even earned the nickname of ”Achilles’ heel of genomes” (Galtier and Duret, 2007). Galtier et al. (2018) proposed that the negative relationship between gBGC and effective population size observed in angiosperms and animals could also arise from a drift-barrier effect, where a low effective population size imposes a limit to the efficacy of selection against gBGC. Here we show that despite a deleterious effect at the population level, gBGC is still (weakly) positively selected. Therefore, the levels of gBGC observed in animals may not be counter-selected at all. In this view, the pervasive existence of gBGC in eukaryotes is not explained by a limited efficiency of negative selection due to drift, but by the short-term advantage of biasing gene conversion towards GC that limits the long-term reproductive capacity of the population as a whole.

### The strength of selection acting on gBGC

The strength of stabilizing selection on gBGC according to our model is weak, and as a consequence, the equilibrium variance can be substantial. Of note, our estimate of the equilibrium variance depends on several assumptions. First, we ignored linkage and only considered the direct conversion gain/cost in fitness in one generation, while in fact, a gBGC modifier will be statistically linked to the deleterious AT alleles it corrects over several generations. By considering only the direct gain/cost at one generation, we might be underestimating the strength of selection acting on a modifier, and thus overestimating the evolutionary variance. However, the agreement of our analytical estimate with the simulations (which do incorporate the effect of linkage) suggests that the impact of linkage is not major.

Second, when computing the selection induced on a modifier, we implicitly assume that the population has time to reach mutation-selection-drift-gBGC equilibrium between each modification of *b* (low mutation rate at modifier loci). When the number and effect sizes of loci that can influence the strength of gBGC are large enough, such that the population does not have time to reach mutation-selection-drift-gBGC equilibrium between two consecutive modifications of *b*, the short-term benefit/cost of converting with a bias *b* is not coupled to its long-term benefit/cost. In this case, one should observe increased oscillations around *b∗*, and thus increased evolutionary variance. This point is confirmed by running the simulator under a higher mutation rate at the modifier locus (Figure S1&2).

Empirically, not much is known about the genetic architecture of gBGC, so the effective mutation rate at the gBGC modifiers is difficult to estimate. Essentially, however, the results obtained here, which show a reasonable match with the simulated variance under linkage and assuming a low mutation rate at the modifier, give the best-case scenario among all possible genetic architectures for gBGC, i.e. the scenario for which stabilizing selection on *b* is tightest. Even so, it remains weak, at least too weak to always guarantee a positive gBGC under empirically reasonable conditions (Table 1). On the other hand, selection against excessively high levels of gBGC might still be efficient, depending on the exact distribution of fitness and dominance effects.

### The penetrance of a somatic repair bias in meiosis

In our derivation, we have focussed on the consequences of biased repair in the germ line, disregarding all considerations about somatic constraints. In reality, however, it is very likely that the mechanisms that bias DNA repair towards GC in meiosis are the same as those that operate in somatic cells. Most single nucleotide DNA damages involve wrongly incorporated As, Ts, or even Us. Repair enzymes that minimize the somatic mutation rate should therefore be GC-biased. In this sense, in mammals, the base excision repair pathway has DNA glycosilases for excising adenines and thymines, but none for guanine or cytosine (Krokan and Bjørås, 2013).

Accounting for these somatic constraints leads to a different perspective on the evolution of gBGC, which could then be seen as an indirect consequence of the shared repair machinery between meiosis and somatic repair. In this context, modifiers can still act on the strength of gBGC, although now by modulating the penetrance of the structurally GC-biased repair system in meiosis. The simulation model could easily be adapted to incorporate such somatic constraints, essentially by assuming that the modifiers of gBGC would act multiplicatively on *b*, itself a priori assumed positive. The semi-analytical derivation is less dependent on such details, as it merely quantifies the selection induced at equilibrium on any modifier of gBGC.

These considerations lead to a re-interpretation of the results presented here. What they fundamentally suggest is that, depending on the exact distribution of fitness and dominance effects, there might be enough selection for limiting the penetrance of the somatic repair bias in meiosis, if this ever leads to overly strong gBGC (deleterious at the individual level). In this view, the expected strength of gBGC should lie between the somatic repair bias (strongly GC biased) and *b*^*∗*^. This could explain why meiotic gene conversion appears to be universally GC-biased, despite the weak selection preventing it from being AT-biased in low *N*_*e*_ species (Table 1). Of note, even if selection to limit the penetrance of the somatic repair bias is maximally effective, the expected value of *b* still induces a substantial load at the population level.

### gBGC and effective population size

In mammals, there is a (weak) correlation between effective population size *N*_*e*_ and the population-scaled gBGC coefficient (*B* = 4*N*_*e*_*b*) (Lartillot, 2013; Galtier, 2021). This correlation is also observed among human populations (Glémin et al., 2015; Subramanian, 2019), and effective population size seems to explain the difference in *B* between two passerine species (Barton and Zeng, 2021). However, in *Leptidea* butterflies there is no relationship between *B* and genetic diversity, suggesting that the transmission bias *b* is lower in species/populations of higher effective population size (Boman et al., 2021). Finally, no correlation has been observed between *B* and *N*_*e*_ in 29 animal species (Galtier et al., 2018), or in 11 species of angiosperms (Clément et al., 2017).

The most probable hypothesis so far is that in animals and plants, there is a negative correlation between the repair bias *b*_0_ and effective population size (Galtier et al., 2018). Several arguments have been proposed to explain this negative relationship. As previously said, Galtier et al. (2018) proposed a drift-barrier hypothesis: assuming that gBGC is deleterious, it can be more efficiently counter-selected in species with higher *N*_*e*_. Our modeling work provides another interpretation, by pointing out that the evolutionary optimum *b*^*∗*^ is itself negatively correlated with *N*_*e*_.

On the other hand, it has been shown that depending on the repair mechanisms, the intensity of the bias could be negatively correlated with heterozygosity (Lesecque et al., 2013; Li et al., 2019). As heterozygosity is supposed to be proportional to *N*_*e*_, this can also explain why we observe no correlation between *B* and *N*_*e*_ (Clément et al., 2017; Galtier et al., 2018; Boman et al., 2021), while we still expect one under selection only. However, the switch to such heterozygosity-dependent mechanisms could also be an adaptive response to the increasing cost of gBGC, and the two hypotheses are not mutually exclusive. Nevertheless, these hypothesis remain verbal, and a proper modelling of the molecular mechanisms of gBGC and their selective advantage is needed to put them to the test.

### Empirical relevance

We highlighted that gBGC being deleterious at the population level is not an indicator that it is negatively selected. It is therefore unclear whether the levels of gBGC currently found in eukaryotes are actually negatively selected. Here, we computed the expected *b*^*∗*^ under empirically realistic parameters, and recover a rather high *b*^*∗*^. Of note, in the context of a heterogeneous recombination landscape, such as considered above, most of the selection induced on the modifier is contributed by those positions that are under the strongest gBGC, which correspond to the highly recombining regions of the genome. Our model therefore predicts even higher equilibrium values for *b*^*∗*^ under more homogeneous recombination landscapes (Table S1).

It is important to note, however, that this estimation is sensitive to parameters that are very difficult to estimate reliably. Notably, the size of the genome that is under selection(Rands et al., 2014), the DFE and more specifically the size of the compartment of strongly deleterious mutations. Moreover, *b*^*∗*^ is strongly sensitive to the distribution of dominance effects, about which little is known (Billiard et al., 2021). Finally, it relies on the assumption that half the genome under selection has a GC allele as optimal and the other half an AT allele. This assumption is intuitive and is made in almost all models of gBGC (Bengtsson, 1990; Glémin, 2010; Bolívar et al., 2016; Corcoran et al., 2017). However, when using empirical fitness landscapes in protein coding genes instead of an arbitrary distribution of selective effects, it appears that AT encoded amino-acids are more often optimal than GC-ones, which seems to be partly due to the structure of the genetic code (Joseph, 2024). Under this scenario, a slight mutational bias is actually beneficial, and thus *b*^*∗*^ should be lower.

Overall, while the present model significantly improves our understanding of the selective pressures exerted on gBGC, it is by no mean an attempt to accurately predict the strength of gBGC *in natura*.

## 4 Methods

### 4.1 Model

The model assumes a population of fixed size *N* diploid individuals, randomly mating and with non-overlapping generations. The genome is composed of a single chromosome. Since neutral loci don’t have any impact on the evolution of gBGC, they are not explicitly modeled. As a result, the chromosome is assumed to consist of *L* bi-allelic positions, with two alternative alleles, *W* (weak) or *S* (strong), that are all under selection with locus-specific selective strengths. The model also invokes a modifier locus placed somewhere along the chromosome (in the experiments conducted here, at one third of the total length of the chromosome).

For a given selected position *i*, 1 *≤ i ≤ L*, either the *W* allele or the *S* allele is considered deleterious with probability 1*/*2, in which case the selection strength *s*_*i*_ acting on the deleterious allele is randomly drawn from a gamma distribution of mean 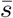 and shape parameter *a*. All selected loci share the same dominance coefficient *h*. In the following, the co-dominant case *h* = 1*/*2 and the recessive case, where the deleterious allele is recessive with 0 *< h <* 1*/*2, are both considered. The selective effects are assumed additive over loci. Thus, assuming locus *i* is such that *W* is the deleterious allele, then the log-fitness contribution is 0 for genotype *SS, hs*_*i*_ for genotype *SW* and *s*_*i*_ for genotype *WW* (and conversely for loci for which *S* is the deleterious allele). Letting 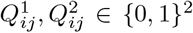 stand for the genotype of diploid individual *j* at position *i*, with the convention that 1 stands for the deleterious allele (which can be either *S* or *W* depending on the locus), the total Malthusian (log) fitness of individual *j* is then given by:

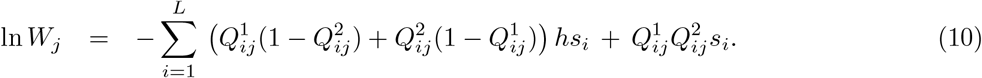

The selected positions undergo recurrent mutations between *W* and *S*. Allele *W* mutates towards *S* at rate *u*, and allele *S* mutates towards *W* at rate *λu* per generation.

The modifier locus encodes an additive quantitative trait controlling the bias of gene conversion. For individual *j*, with genotype 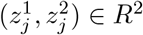 at the modifier locus, the bias is then equal to:

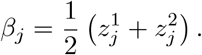

How this bias exactly impacts gene conversion during meiosis is described below. The modifier locus mutates are rate *w*, in which case the quantitative contribution of the mutant allele is equal to that of its parent, plus a normally distributed increment, of mean 0 and standard deviation Δ*z*:

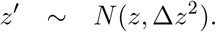

A simplified version of meiosis is implemented as follows. Consider individual *j*. First, each selected position that happens to be heterozygous in this individual undergoes gene conversion with probability *α*, in which case conversion is towards the *S* allele with probability (1 + *β*_*j*_)*/*2 and towards the *W* allele with probability (1 *− β*_*j*_)*/*2, with *β*_*j*_ such as defined above (equation..). Second, a cross-over point is chosen uniformly at random over the chromosome, and two recombinant chromosomes are produced by swapping the segments on both sides of the cross-over point. Thus, both the rate of cross-over and the rate of gene conversion are considered fixed and invariant across individuals, while the bias of the gene conversion events is allowed to vary between individuals, based on the genotype at the modifying locus. Of note, a positive (resp. negative) value for *β*_*j*_ results in biased gene conversion towards *S* (resp. towards *W*). Quantitatively, at a given selected position at which individual *j* is heterozygous, the net proportions of gametes produced by this individual bearing the *S* allele is:

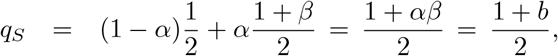

with *b* = *αβ*. Similarly, the proportion of gametes with the *W* allele is 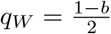.

The overall life cycle runs as follows. First, all individuals of the current generation undergo mutations both at the modifying and at the selected loci, with mutation rates such as given above. Next, each individual of the next generation is produced by first randomly choosing two parents in the current generation, each with a probability proportional to is fitness *W* (such as given by equation 10 above). Each of the two chosen individuals then undergoes a meiosis, producing a pair of gametes, one of which is randomly picked out and paired with the gamete produced by applying the same procedure to the other individual. Of note, only one gamete per meiosis is kept for the next generation, the other one being discarded.

Altogether, the parameters of the model are:

*N* : population size

*L*: number of loci under selection

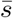: mean selection strength at the selected loci

*a*: shape of the distribution of selection strengths across loci

*h*: dominance coefficient

*u*: basal mutation rate at the selected loci

*λ*: mutation bias (*S ↦ W* relative to *W ↦ S*) *w*: mutation rate at the modifier locus

Δ*z*: mean effect size of the mutations at the modifier locus

*α*: gene conversion rate (per generation and per selected locus)

### 4.2 Theory / Analytical approximation

Here, a semi-analytical approximation is derived for determining the equilibrium value of the net strength of biased gene conversion *b* as well as its evolutionary variance. This derivation assumes a low mutation rate at the modifier locus (low *w*), such that the population is, at any time, approximately monomorphic for the strength of gBGC, and all selected loci are at mutation-selection-drift-conversion equilibrium under this value of gBGC. The derivation also assumes that linkage both among selected loci and between the modifier and the selected loci is negligible. The first condition implies that background selection is weak, and that the mutation-selection-drift-conversion equilibrium can be determined independently at each locus. The second is motivated by the fact that, in practice, most selected loci are sufficiently far from the modifier, such that most of the induced selection is contributed by loci that not tightly linked with the modifier.

Consider a population monomorphic at the modifier locus for an allele of strength *β*, at equilibrium under a gBGC of strength *b* = *αβ*. In this population, a mutant at the modifier locus, of size 2*δβ* appears in an individual. This individual thus has a gBGC strength of *b*^*′*^ = *b* + *δb* in its germline, with *δb* = *αδβ*. We want to determine the net selective advantage or disadvantage incurred by this individual, owing to its departure from the population-level gBGC. This selection will be indirectly contributed by the effect of biased gene conversion on the selected loci across the genome. Therefore, in the following, this will be called the selection *induced* on the gBGC modifier, or induced selection for short.

Under efficient linkage dissipation, induced selection is the sum of the contributions of all selected loci. Consider in a first step a single locus at which *W* is the deleterious allele, with selection *s*, dominance *h* and segregating in the population at frequency *x*. Given *x*, the probability for the individual to be heterozygous at this position is:

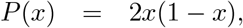

in which case the S and W allele are transmitted in the gametes with probability 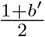 and 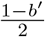 *′*, respectively. In a random mating population, this will result in an average fitness gain in the offspring of:

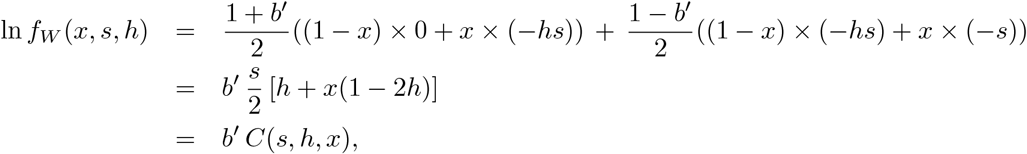

where:

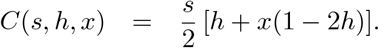

Of note, if *b*^*′*^ *>* 0, this is indeed a gain, since on average, *S* alleles, which have a higher fitness at that position, are over-transmitted. Next, to assess the fate of the gBGC mutant, one should discount the equivalent gain, but under a gBGC equal to *b* in the population, such that the average selective advantage contributed by the selected position under consideration to the individual bearing the mutant allele for the modifier (now accounting for the probability for this individual to be a heterozygote at the selected locus) is:

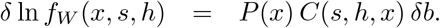

Equation 11 gives the cost conditional on the frequency *x* of the *W* allele at the focal selected position and conditional on the selection coefficient *s*. This needs to be averaged over *x* at mutation-selection-conversion-drift equilibrium (here noted 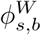) and then summed over the distribution of selective effects across the *L/*2 loci being deleterious towards the *W* allele:

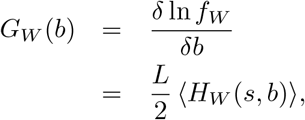

where the angle brackets stand for the expectation over *s* under the DFE, and,

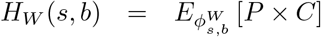

is the expectation over *x* under 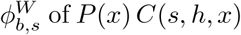 In other words, it is the net gain induced by conversion events at loci that are *W* -deleterious, with selection coefficient *s* and dominance coefficient *h*. In turn, the distribution 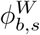 is given by (Wright, Glemin):

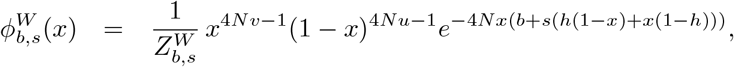

where 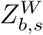 is the normalization constant:

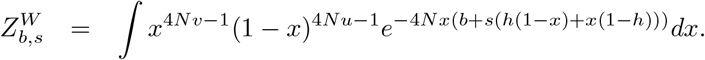

A similar derivation is done for a locus at which *W* is the deleterious allele, which, by symmetry, gives:

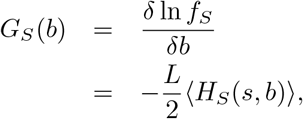

where

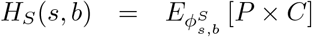

and

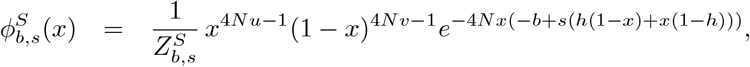

with normalization constant:

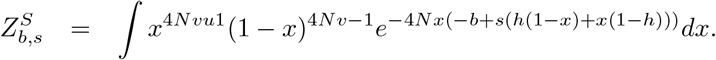

Of note, *P* (*x*) and *C*(*s, h, x*) are positive for all *x*, and thus, increasing biased gene conversion towards the strong alleles results in a net gain over *W* -deleterious loci, but a net a loss over *S*-deleterious loci. Whether the mutant for gBGC is favoured by this induced selection will depend on the balance between these two components. In other words, the total selection induced on the modifier is:

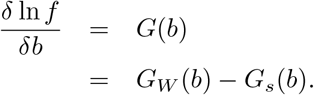

### 4.3 Implementation

The model was implemented in C++, using openMP for parallelizing the computations. For all results presented here, it was run under population sizes of size *N* = 1000, with *L* = 10000 selected loci, for a total of 210 000 generations, discarding the first 10 000 generations (burn-in) and subsampling 1 every 100 generations, upon which averages and standard deviations for quantities of interest were computed on the remaining 2000 points.

Numerical integration and solving was done in Python, using the scipy library for numerical integration over the allele frequency distributions. For integrating over the gamma distribution of selective effects, the gamma distribution was discretized into *n* = 300 points, corresponding to the mid-points of the successive 1*/n* quantiles, and then the integral over the distribution was approximated as the equal-weighted average of the integrand over these *n* values for *hs*.

## Acknowledgments

We are grateful to Sylvain Glémin, Laurent Duret and Nicolas Galtier for their comments on previous versions of this manuscript. We also would like to thank Sylvain Glémin for insightful discussions on the subject and on the model.

## Data and materials availability

All the scripts necessary to reproduce this study are available at https://gitlab.in2p3.fr/bayesiancook/gbgc

## References

Barton, H. J. and Zeng, K. (2021). The effective population size modulates the strength of GC biased gene conversion in two passerines.

Bengtsson, B. O. (1986). Biased conversion as the primary function of recombination. Genetics Research, 47(1):77–80.

Bengtsson, B. O. (1990). The effect of biased conversion on the mutation load. Genetics Research, 55(3):183– 187.

Bengtsson, B. O. and Uyenoyama, M. K. (1990). Evolution of the segregation ratio: Modification of gene conversion and meiotic drive. Theoretical Population Biology, 38(2):192–218.

Berglund, J., Pollard, K. S., and Webster, M. T. (2009). Hotspots of Biased Nucleotide Substitutions in Human Genes. PLOS Biology, 7(1):e1000026.

Billiard, S., Castric, V., and Llaurens, V. (2021). The integrative biology of genetic dominance. Biological Reviews, 96(6):2925–2942.

Bolívar, P., Guéguen, L., Duret, L., Ellegren, H., and Mugal, C. F. (2019). GC-biased gene conversion conceals the prediction of the nearly neutral theory in avian genomes. Genome Biol, 20(1):5.

Bolívar, P., Mugal, C. F., Nater, A., and Ellegren, H. (2016). Recombination Rate Variation Modulates Gene Sequence Evolution Mainly via GC-Biased Gene Conversion, Not Hill–Robertson Interference, in an Avian System. Mol Biol Evol, 33(1):216–227.

Boman, J., Mugal, C. F., and Backström, N. (2021). The Effects of GC-Biased Gene Conversion on Patterns of Genetic Diversity among and across Butterfly Genomes. Genome Biology and Evolution, 13(5).

Brown, T. C. and Jiricny, J. (1987). A specific mismatch repair event protects mammalian cells from loss of 5-methylcytosine. Cell, 50(6):945–950.

Castellano, D., Maciá, M. C., Tataru, P., Bataillon, T., and Munch, K. (2019). Comparison of the Full Distribution of Fitness Effects of New Amino Acid Mutations Across Great Apes. Genetics, 213(3):953– 966.

Clément, Y., Sarah, G., Holtz, Y., Homa, F., Pointet, S., Contreras, S., Nabholz, B., Sabot, F., Sauné, L., Ardisson, M., Bacilieri, R., Besnard, G., Berger, A., Cardi, C., Bellis, F. D., Fouet, O., Jourda, C., Khadari, B., Lanaud, C., Leroy, T., Pot, D., Sauvage, C., Scarcelli, N., Tregear, J., Vigouroux, Y., Yahiaoui, N., Ruiz, M., Santoni, S., Labouisse, J.-P., Pham, J.-L., David, J., and Glémin, S. (2017). Evolutionary forces affecting synonymous variations in plant genomes. PLOS Genetics, 13(5):e1006799.

Corcoran, P., Gossmann, T. I., Barton, H. J., The Great Tit HapMap Consortium, Slate, J., and Zeng, K. (2017). Determinants of the Efficacy of Natural Selection on Coding and Noncoding Variability in Two Passerine Species. Genome Biology and Evolution, 9(11):2987–3007.

Duret, L. and Arndt, P. F. (2008). The Impact of Recombination on Nucleotide Substitutions in the Human Genome. PLOS Genetics, 4(5):e1000071.

Duret, L. and Galtier, N. (2009). Biased Gene Conversion and the Evolution of Mammalian Genomic Landscapes. Annu. Rev. Genom. Hum. Genet., 10(1):285–311.

Eyre-Walker, A. (1999). Evidence of Selection on Silent Site Base Composition in Mammals: Potential Implications for the Evolution of Isochores and Junk DNA. Genetics, 152(2):675–683.

Eyre-Walker, A. and Keightley, P. D. (2007). The distribution of fitness effects of new mutations. Nat Rev Genet, 8(8):610–618.

Eyre-Walker, A., Woolfit, M., and Phelps, T. (2006). The Distribution of Fitness Effects of New Deleterious Amino Acid Mutations in Humans. Genetics, 173(2):891–900.

Galtier, N. (2021). Fine-scale quantification of GC-biased gene conversion intensity in mammals. Peer Community Journal, 1.

Galtier, N. and Duret, L. (2007). Adaptation or biased gene conversion? Extending the null hypothesis of molecular evolution. Trends in Genetics, 23(6):273–277.

Galtier, N., Duret, L., Glémin, S., and Ranwez, V. (2009). GC-biased gene conversion promotes the fixation of deleterious amino acid changes in primates. Trends in Genetics, 25(1):1–5.

Galtier, N., Piganeau, G., Mouchiroud, D., and Duret, L. (2001). GC-Content Evolution in Mammalian Genomes: The Biased Gene Conversion Hypothesis. Genetics, 159(2):907–911.

Galtier, N., Roux, C., Rousselle, M., Romiguier, J., Figuet, E., Glémin, S., Bierne, N., and Duret, L. (2018). Codon Usage Bias in Animals: Disentangling the Effects of Natural Selection, Effective Population Size, and GC-Biased Gene Conversion. Molecular Biology and Evolution, 35(5):1092–1103.

Glémin, S. (2010). Surprising Fitness Consequences of GC-Biased Gene Conversion: I. Mutation Load and Inbreeding Depression. Genetics, 185(3):939–959.

Glémin, S., Arndt, P. F., Messer, P. W., Petrov, D., Galtier, N., and Duret, L. (2015). Quantification of GC-biased gene conversion in the human genome. Genome Res., 25(8):1215–1228.

Joseph, J. (2024). Increased Positive Selection in Highly Recombining Genes Does not Necessarily Reflect an Evolutionary Advantage of Recombination. Molecular Biology and Evolution, 41(6):msae107.

Kostka, D., Hubisz, M. J., Siepel, A., and Pollard, K. S. (2012). The Role of GC-Biased Gene Conversion in Shaping the Fastest Evolving Regions of the Human Genome. Molecular Biology and Evolution, 29(3):1047–1057.

Krokan, H. E. and Bjørås, M. (2013). Base Excision Repair. Cold Spring Harb Perspect Biol, 5(4):a012583.

Lachance, J. and Tishkoff, S. A. (2014). Biased Gene Conversion Skews Allele Frequencies in Human Populations, Increasing the Disease Burden of Recessive Alleles. The American Journal of Human Genetics, 95(4):408–420.

Lartillot, N. (2013). Phylogenetic Patterns of GC-Biased Gene Conversion in Placental Mammals and the Evolutionary Dynamics of Recombination Landscapes. Molecular Biology and Evolution, 30(3):489–502.

Lesecque, Y., Mouchiroud, D., and Duret, L. (2013). GC-Biased Gene Conversion in Yeast Is Specifically Associated with Crossovers: Molecular Mechanisms and Evolutionary Significance. Molecular Biology and Evolution, 30(6):1409–1419.

Li, R., Bitoun, E., Altemose, N., Davies, R. W., Davies, B., and Myers, S. R. (2019). A high-resolution map of non-crossover events reveals impacts of genetic diversity on mammalian meiotic recombination. Nat Commun, 10(1):3900.

Long, H., Sung, W., Kucukyildirim, S., Williams, E., Miller, S. F., Guo, W., Patterson, C., Gregory, C., Strauss, C., Stone, C., Berne, C., Kysela, D., Shoemaker, W. R., Muscarella, M. E., Luo, H., Lennon, J. T., Brun, Y. V., and Lynch, M. (2018). Evolutionary determinants of genome-wide nucleotide composition. Nat Ecol Evol, 2(2):237–240.

Mancera, E., Bourgon, R., Brozzi, A., Huber, W., and Steinmetz, L. M. (2008). High-resolution mapping of meiotic crossovers and non-crossovers in yeast. Nature, 454(7203):479–485.

Meunier, J. and Duret, L. (2004). Recombination Drives the Evolution of GC-Content in the Human Genome. Molecular Biology and Evolution, 21(6):984–990.

Nagylaki, T. (1983). Evolution of a finite population under gene conversion. Proceedings of the National Academy of Sciences, 80(20):6278–6281.

Necşulea, A., Popa, A., Cooper, D. N., Stenson, P. D., Mouchiroud, D., Gautier, C., and Duret, L. (2011). Meiotic recombination favors the spreading of deleterious mutations in human populations. Human Mutation, 32(2):198–206.

Ohta, T. (1992). The Nearly Neutral Theory of Molecular Evolution. Annual Review of Ecology and Systematics, 23:263–286.

Pessia, E., Popa, A., Mousset, S., Rezvoy, C., Duret, L., and Marais, G. A. (2012). Evidence for widespread GC-biased gene conversion in eukaryotes. Genome biology and evolution, 4(7):675–682.

Pouyet, F., Aeschbacher, S., Thiéry, A., and Excoffier, L. (2018). Background selection and biased gene conversion affect more than 95% of the human genome and bias demographic inferences. eLife, 7:e36317.

Pouyet, F., Mouchiroud, D., Duret, L., and Sémon, M. (2017). Recombination, meiotic expression and human codon usage. eLife, 6:e27344.

Pratto, F., Brick, K., Khil, P., Smagulova, F., Petukhova, G. V., and Camerini-Otero, R. D. (2014). Recombination initiation maps of individual human genomes. Science, 346(6211):1256442–1256442.

Rands, C. M., Meader, S., Ponting, C. P., and Lunter, G. (2014). 8.2% of the Human Genome Is Constrained: Variation in Rates of Turnover across Functional Element Classes in the Human Lineage. PLOS Genetics, 10(7):e1004525.

Ratnakumar, A., Mousset, S., Glémin, S., Berglund, J., Galtier, N., Duret, L., and Webster, M. T. (2010). Detecting positive selection within genomes: the problem of biased gene conversion. Philosophical Transactions of the Royal Society B: Biological Sciences, 365(1552):2571–2580.

Roman, H. (1985). Gene conversion and crossing-over. Environmental Mutagenesis, 7(6):923–932.

Smagulova, F., Gregoretti, I. V., Brick, K., Khil, P., Camerini-Otero, R. D., and Petukhova, G. V. (2011). Genome-wide analysis reveals novel molecular features of mouse recombination hotspots. Nature, 472(7343):375–378.

Smeds, L., Mugal, C. F., Qvarnström, A., and Ellegren, H. (2016). High-Resolution Mapping of Crossover and Non-crossover Recombination Events by Whole-Genome Re-sequencing of an Avian Pedigree. PLOS Genetics, 12(5):e1006044.

Subramanian, S. (2019). Population size influences the type of nucleotide variations in humans. BMC Genet, 20(1):1–12.

Webster, M. T., Axelsson, E., and Ellegren, H. (2006). Strong Regional Biases in Nucleotide Substitution in the Chicken Genome. Molecular Biology and Evolution, 23(6):1203–1216.

Winkler, H. (1930). Die Konversion der Gene : eine vererbungstheoretische Untersuchung. G Fischer.

