## Supplementary Material for "The evolution of GC-biased gene conversion by means of natural selection"

### Supplementary materials from: The evolution of GC-biased gene conversion by means of natural selection

June 21, 2024

| | | $h\bar{s} = 0.01$ | | | $h\bar{s} = 0.05$ | | |
|---|---|---|---|---|---|---|---|
| $h$ | $N$ | $B_h^*$ | stdev | $p(B < 0)$ | $B_h^*$ | stdev | $p(B < 0)$ |
| 0.5 | $10^4$ | 11.9 | 2.5 | $< 0.001$ | 25.4 | 3.8 | $< 0.001$ |
| | $10^5$ | 36.7 | 4.6 | $< 0.001$ | 89.9 | 3.4 | $< 0.001$ |
| | $10^6$ | 157.7 | 3.4 | $< 0.001$ | 567.3 | 3.6 | $< 0.001$ |
| 0.1 | $10^4$ | 6.9 | 1.3 | $< 0.001$ | 9.3 | 1.3 | $< 0.001$ |
| | $10^5$ | 10.8 | 1.3 | $< 0.001$ | 13.3 | 1.3 | $< 0.001$ |
| | $10^6$ | 23.9 | 2.1 | $< 0.001$ | 31.3 | 2.4 | $< 0.001$ |
| 50% 0.1 : 50% 0.5 | $10^4$ | 8.0 | 1.5 | $< 0.001$ | 11.1 | 1.5 | $< 0.001$ |
| | $10^5$ | 13.1 | 1.5 | $< 0.001$ | 16.3 | 1.5 | $< 0.001$ |
| | $10^6$ | 32.3 | 2.7 | $< 0.001$ | 44.6 | 3.2 | $< 0.001$ |
| 10% 0.1 : 90% 0.5 | $10^4$ | 10.2 | 2.0 | $< 0.001$ | 15.7 | 1.9 | $< 0.001$ |
| | $10^5$ | 19.6 | 2.1 | $< 0.001$ | 26.1 | 2.2 | $< 0.001$ |
| | $10^6$ | 64.5 | 3.9 | $< 0.001$ | 94.2 | 4.7 | $< 0.001$ |

Table S1: Numerical estimates of scaled intensity of gBGC $B^* = 4Nb^*$ (mean over the genome), equilibrium standard deviation and probability of a negative gBGC, for different parameter values for $N$, $h$, $h\bar{s}$, with a homogeneous recombination landscape.

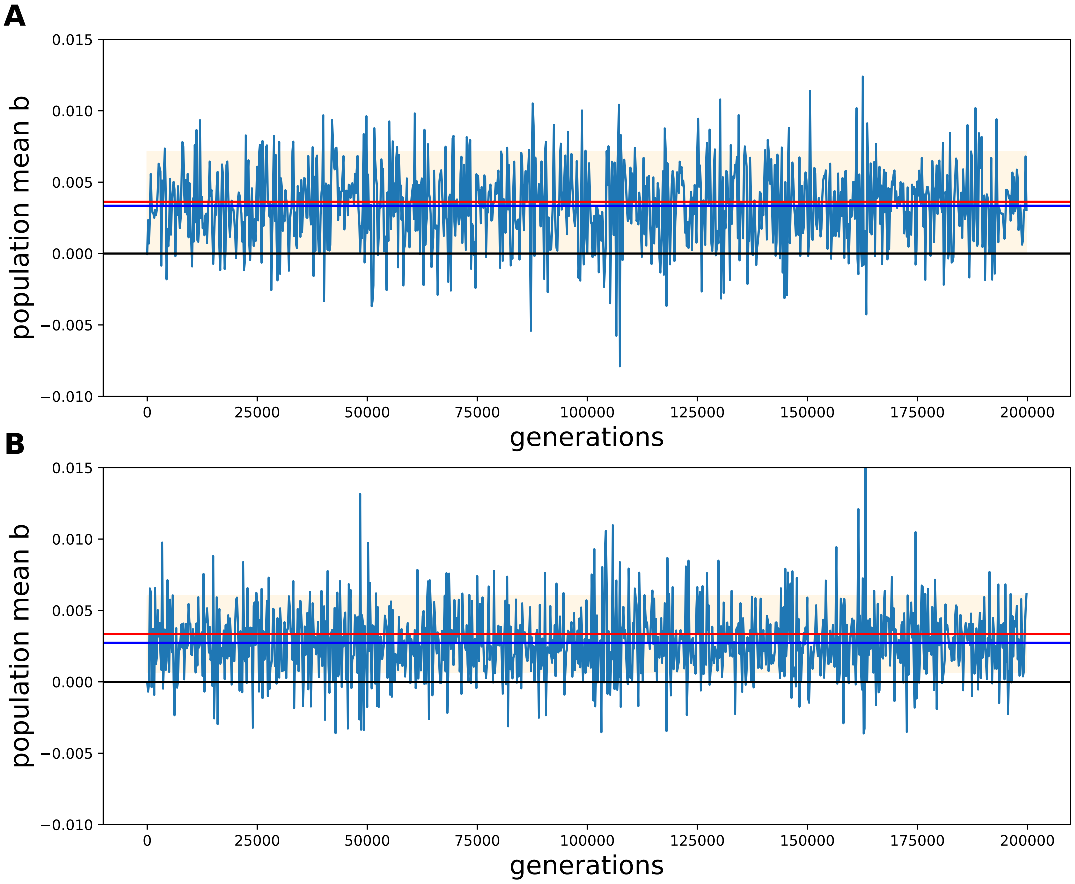


Figure S1: Evolution of the strength of gBGC (mean of $b$ over the population) over the generations, for the co-dominant (A) and recessive (B) cases under under a mutation rate at the modifier loci of $w = 10^{-3}$ ($Nw = 1$). Blue horizontal line: mean over the entire run; red horizontal line: equilibrium value predicted by the analytical approximation; shaded area: predicted equilibrium variance.

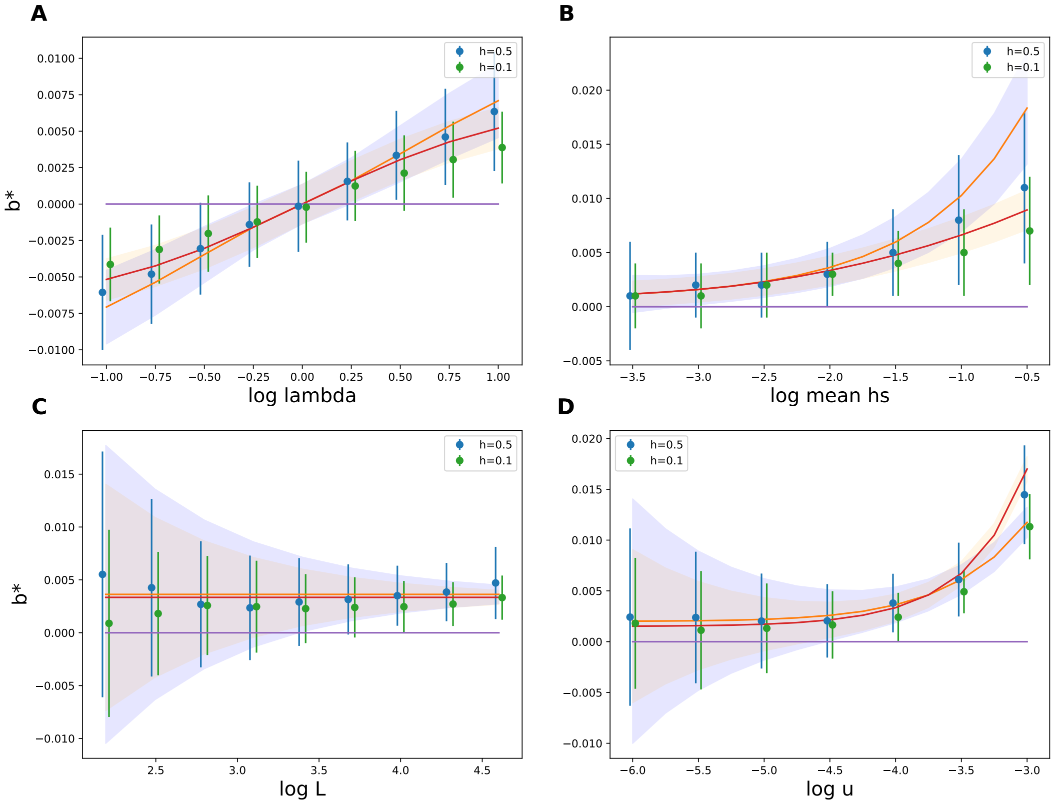


Figure S2: Mean equilibrium $b^*$ and standard deviation, as a function of mutation bias $\lambda$ (A), mean selective effect $h\bar{s}$ (B), number of selected positions in the genome $L$ (C) and mutation rate $u$ (D), under the co-dominant (blue) and the recessive (orange) case, obtained by simulations (dots and associated vertical bars) and predicted by the analytical approximation (curve and associated shaded area), under a mutation rate at the modifier loci of $w = 10^{-3}$ ($Nw = 1$).

| | | $h\bar{s} = 0.01$ | | | $h\bar{s} = 0.05$ | | |
|---|---|---|---|---|---|---|---|
| $h$ | $N$ | $B_h^*$ | stdev | $p(B < 0)$ | $B_h^*$ | stdev | $p(B < 0)$ |
| | $10^4$ | 1.3 | 1.1 | 0.108 | 2.5 | 1.5 | 0.049 |
| 0.5 | $10^5$ | 3.4 | 1.8 | 0.028 | 6.8 | 1.8 | $< 0.001$ |
| | $10^6$ | 10.9 | 1.4 | $< 0.001$ | 21.6 | 0.7 | $< 0.001$ |
| | $10^4$ | 0.7 | 0.5 | 0.083 | 0.8 | 0.5 | 0.041 |
| 0.1 | $10^5$ | 0.9 | 0.5 | 0.027 | 1.1 | 0.5 | 0.010 |
| | $10^6$ | 1.8 | 0.7 | 0.004 | 2.2 | 0.7 | 0.001 |
| | $10^4$ | 0.8 | 0.6 | 0.081 | 1.0 | 0.6 | 0.033 |
| 50% 0.1 : 50% 0.5 | $10^5$ | 1.2 | 0.6 | 0.021 | 1.4 | 0.6 | 0.006 |
| | $10^6$ | 2.5 | 0.9 | 0.003 | 3.1 | 0.9 | $< 0.001$ |
| | $10^4$ | 1.1 | 0.8 | 0.086 | 1.5 | 0.8 | 0.024 |
| 10% 0.1 : 90% 0.5 | $10^5$ | 1.8 | 0.8 | 0.012 | 2.3 | 0.8 | 0.002 |
| | $10^6$ | 4.8 | 1.6 | 0.001 | 6.7 | 1.5 | $< 0.001$ |

Table S2: Numerical estimates of scaled intensity of gBGC $B^* = 4Nb^*$ (mean over the genome), equilibrium standard deviation and probability of a negative gBGC, for different parameter values for $N$, $h$, $h\bar{s}$, with a shape parameter $a = 0.1$ for the distribution of fitness effects.

| | | $h\bar{s} = 0.01$ | | | $h\bar{s} = 0.05$ | | |
|---|---|---|---|---|---|---|---|
| $h$ | $N$ | $B_h^*$ | stdev | $p(B < 0)$ | $B_h^*$ | stdev | $p(B < 0)$ |
| | $10^4$ | 1.7 | 0.9 | 0.024 | 4.3 | 1.4 | 0.001 |
| 0.5 | $10^5$ | 6.7 | 1.6 | $< 0.001$ | 19.2 | 0.7 | $< 0.001$ |
| | $10^6$ | 38.4 | 0.9 | $< 0.001$ | 124.0 | 0.8 | $< 0.001$ |
| | $10^4$ | 1.0 | 0.4 | 0.009 | 1.3 | 0.4 | $< 0.001$ |
| 0.1 | $10^5$ | 1.6 | 0.4 | $< 0.001$ | 2.0 | 0.4 | $< 0.001$ |
| | $10^6$ | 4.2 | 0.8 | $< 0.001$ | 6.0 | 1.0 | $< 0.001$ |
| | $10^4$ | 1.1 | 0.5 | 0.009 | 1.5 | 0.4 | $< 0.001$ |
| 50% 0.1 : 50% 0.5 | $10^5$ | 1.9 | 0.5 | $< 0.001$ | 2.4 | 0.5 | $< 0.001$ |
| | $10^6$ | 5.8 | 1.1 | $< 0.001$ | 8.8 | 1.1 | $< 0.001$ |
| | $10^4$ | 1.4 | 0.6 | 0.012 | 2.2 | 0.6 | $< 0.001$ |
| 10% 0.1 : 90% 0.5 | $10^5$ | 2.9 | 0.7 | $< 0.001$ | 4.1 | 0.8 | $< 0.001$ |
| | $10^6$ | 10.7 | 1.0 | $< 0.001$ | 15.3 | 0.9 | $< 0.001$ |

Table S3: Numerical estimates of scaled intensity of gBGC $B^* = 4Nb^*$ (mean over the genome), equilibrium standard deviation and probability of a negative gBGC, for different parameter values for $N$, $h$, $h\bar{s}$, , with a shape parameter $a = 0.3$ for the distribution of fitness effects.
